# BMP signaling modulations control primitive streak patterning

**DOI:** 10.1101/2024.10.01.616050

**Authors:** Gaël Simon, Jean-Louis Plouhinec, Pascale Gilardi-Hebenstreit, Benoit Sorre, Jérôme Collignon

## Abstract

The primitive streak (PS) is the embryonic structure from which mesoderm and definitive endoderm cells emerge to form organs and fetal tissues. The identities of the cells leaving the PS depend on both the time and the position at which they exit. Three signaling molecules, BMP4, WNT3 and NODAL, play key roles in determining these identities, but it has been difficult to disentangle the contribution of each to this process in the mouse embryo. Using 2D gastruloids that recapitulate early aspects of mouse gastrulation *in vitro*, we have previously shown that WNT3 and NODAL promote distal PS fates on their own, and proximal PS fates in the presence of BMP4. We now detail further the impact of BMP4 on the cell identities produced in gastruloids. Single cell RNA-seq analysis shows that prolonged exposure to BMP4 results in the presence of all the cell types, including primordial germ cells, which normally form from the most proximal part of the pluripotent epiblast. We then show that reducing the duration or intensity of gastruloid exposure to BMP4 can allow the emergence of more distal cell identities, such as paraxial mesoderm, alongside anterior PS derivatives. These results, which are consistent with embryological studies, provide key elements for understanding how the varied output of the PS is controlled.

## Introduction

The formation of the primitive streak (PS) marks the beginning of gastrulation in all amniote embryos. In the mouse, the PS appears at the edge of the cup-shaped pluripotent epiblast (Epi), at a position corresponding to the posterior side of the embryo, and extends gradually towards its distal tip. Cells emerging early on from the proximal/posterior region of the PS will migrate proximally to form extra-embryonic mesoderm or laterally to form early embryonic mesoderm derivatives, whereas cells emerging from its distal/anterior region will form axial mesoderm and definitive endoderm (Kinder et al., 1999; Lawson, 1999). Analyses of mutant phenotypes have shown that the signaling molecules WNT3, BMP4 and NODAL, are key players in the specification of the cell identities that exit the PS (Arnold & Robertson, 2009; Brennan et al., 2001; Conlon et al., 1994; Liu et al., 1999; Tam & Loebel, 2007; Winnier et al., 1995). Although there is evidence to suggest that WNT3, BMP4 and NODAL play different roles in this process, it has been difficult to establish what each contributes and how their actions combine.

One difficulty stems from the interdependence of *Wnt3*, *Bmp4* and *Nodal* expressions, another from their earlier roles in embryo patterning (Camacho-Aguilar & Warmflash, 2020; Morgani & Hadjantonakis, 2020). However, an *in vitro* approach that allows close control of morphogen exposure and signaling activities now makes it possible to overcome these difficulties. Epiblast-like cells (EpiLCs) cultured as a monolayer on embryo-size circular adhesive micropatterns self-organize when exposed to BMP4 or WNT3a (a substitute for WNT3) and give rise to distinct combinations of the three embryonic germ layers, arranged in concentric rings, in an ordered and reproducible sequence (Morgani et al., 2018; Plouhinec et al., 2022). Such colonies recapitulate key aspects of mouse gastrulation and are thus called mouse 2D gastruloids (m2Dgas).

The characterization of the development of m2Dgas stimulated by different cocktails of morphogens showed that BMP4, in addition to promoting proximal cell-fates in m2Dgas, actively suppresses distal PS cell-fates (Morgani et al., 2018; Plouhinec et al., 2022). This is consistent with the fact that in the embryo proximal fates emerge from a region that is continuously exposed to BMP4, whereas distal fates emerge from a region that is more distant from the source of diffusing BMP4 molecules and later shows local expression of BMP antagonists. In m2Dgas, like in the embryo, BMP4 exposure induces *Wnt3* expression, which in turn locally reinforces a pre-existing *Nodal* expression (Ben-Haim et al., 2006; Plouhinec et al., 2022). We found that the WNT and ACTIVIN/NODAL signaling pathways are strongly activated in BMP4-stimulated colonies, and actually required to form proximal PS identities (Plouhinec et al., 2022). This indicates that, although BMP4 prevents NODAL and WNT3a from activating the distal PS development program, it does not block their signaling pathways but rather restrict them to a specific set of targets.

These m2Dgas studies provided a highly contrasted, but incomplete, view of the impact of BMP signaling, as colonies were either exposed to BMP4 for the entire duration of the culture, or not at all. In the embryo, before gastrulation begins, BMP4 expression is restricted to the region of the extra-embryonic ectoderm (ExE) that is abutting the proximal Epi (Lawson et al., 1999). BMP4 expression is then induced in extra-embryonic mesoderm emerging from the PS, but remains absent from the PS as it extends distally and its distal part begins to express BMP antagonists. In addition, BMP8b and BMP2, which can activate the same signaling pathway as BMP4, are expressed in the ExE and in the visceral endoderm (VE), an extraembryonic layer that lines the epiblast (Ying et al., 2000; Ying & Zhao, 2001). Thus, because of their initial position relative to the source of BMP ligands, the possibility that they may subsequently move further away, either because of embryo growth or their own active migration, and their proximity to cells producing BMP antagonists, PS cells can be exposed to very different levels of BMP ligands, and for very different lengths of time. To determine how this affects their choice of cell identity, we examined the effect of reducing BMP4 exposure on cell fate specification in m2Dgas. First, this involved characterizing in more detail the different cell identities emerging in BMP4-treated m2Dgas using single-cell RNA-seq analysis. Second, we analyzed how shorter BMP exposure time, lower BMP concentration and timed inhibition of BMP signaling affect the range of cell identities that form on micropatterned colonies. We found that m2Dgas continuously exposed to BMP4 give rise to all the cell identities that normally arise from the most proximal region of the Epi, from surface ectoderm, to PGCs and blood progenitors. We then found that reducing BMP exposure allows more distal mesodermal identities to emerge in m2Dgas, but that even a short exposure of progenitor cells to BMP4 is sufficient to alter the fate of their progeny.

## Results

### Detailed characterization of the cell-types emerging in BMP4-induced m2Dgas

Mouse 2D gastruloids were produced using the same protocol as before (Simon et al., 2022). Using the 10x Genomics technology, a single-cell RNA-seq analysis was performed on some of these colonies at five time points before and after the start of BMP4 exposure (0h, 6h, 24h, 48h and 72h) (Fig. 1A), covering their transition from EpiLCs to ectodermal and PS derivatives.

**Figure 1.**
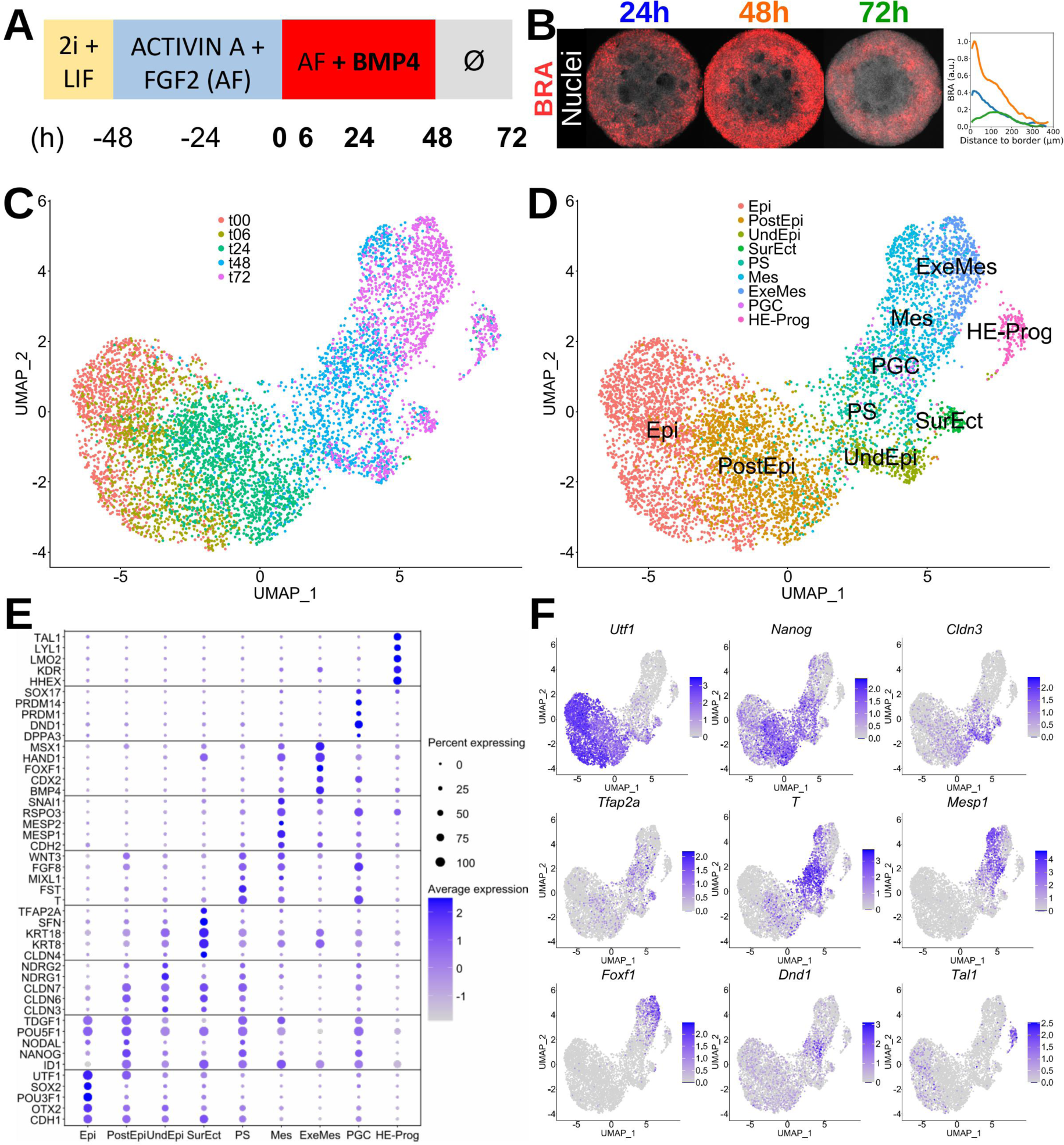
**Single cell RNAseq analysis of the emergence of dis;nct cell iden;;es in BMP4-s;mulated mouse 2D gastruloids** ***(A)*** Culture conditions followed to produce m2Dgas. Cells were seeded on micro-pa9erns at t=-24h. S-mula-on with BMP4 started at t=0h, but from t=48h only basal culture medium was used. Samples were collected for scRNAseq analysis at t=0, 6, 24, 48 and 72h. ***(B)*** Maximum intensity projections of m2Dgas (700 µm) immuno-stained for BRA 24, 48, or 72h aQer the start of BMP4 s-mula-on. ***(C)*** UMAP representation of all the cells analyzed at the different -mes. ***(D)*** UMAP representation of the different cell identity clusters annotated in the dataset. ***(E)*** Dot plot showing the expression of 45 genes across the 9 annotated cell iden--es, and iden-fying 5 genes whose co-expression constitutes a signature for each cell identity. <u>Epi:</u> Epiblast, <u>PostEpi:</u> Posterior Epiblast, <u>UndEpi:</u> Undifferentiated Epiblast, <u>SurEct:</u> Surface ectoderm, <u>PS:</u> Primitive Streak, <u>Mes:</u> Mesoderm, <u>ExeMes:</u> Extra-embryonic mesoderm, <u>PGCs:</u> Primordial Germ Cells, <u>HE-Prog:</u> Haemato-Endothelial Progenitors ***(F)*** UMAP plots of the expression of nine genes, each selected as a marker for a par-cular cluster.

To check that their differentiation had proceeded as expected, some of the colonies were immunostained for a number of markers, including BRACHYURY (BRA), a PS marker, and phosphorylated SMAD1 (pSMAD1), which provides a readout for BMP signaling activity (Fig. 1B, S1A). At t=24h, BRA expression was detected as a thin outer ring. By t=48h this expression had strengthened and the ring had widened, but by t=72h the expression had decreased and was restricted to an inner ring. A gradient of BMP activity was detected from 24h to 72h, with higher activities at the border. These patterns were similar to those described in previous studies (Morgani et al., 2018; Plouhinec et al., 2022). In addition, principal component analysis (PCA) of the expression of 17 differentiation markers confirmed that the m2Dgas produced followed developmental trajectories (Fig. S1B, C) similar to those we characterized previously (Plouhinec et al., 2022).

After a quality control step (see ‘Materials and methods’), the number of cells retained for analysis at the 0h, 6h, 24h, 48h and 72h time points was 1284, 994, 1692, 1333 and 1355 respectively. The data from the various time points were combined using the MNN batch-correction method (Haghverdi et al., 2018) (Fig. 1C). Specific combinations of gene expression identified 9 cell clusters (Fig. 1D). After 24h of BMP stimulation the original EpiLC state (*Utf1, Sox2*, *Pou3f1*, *Otx2*, Cdx1), had been replaced by something similar to posterior epiblast (*Tdgf1, Pou5f1, Nodal, Nanog*, *Id1*). After 48h of stimulation, most cells went through a PS state (*Wnt3*, *Fgf8, Mixl1*, *Fst, T*), subsequently differentiating into mesoderm (*Snai1, Rspo3*, *Mesp2*, *Mesp1*, *Cdh2*), extra-embryonic mesoderm (*Msx1*, *Hand1, Foxf1*, *Cdx2*, *Bmp4*) or haemato-endothelial progenitor (*Tal1*, *Lyl1*, *Lmo2*, *Kdr*, *Hhex*) identities. Cells that didn’t go through the PS state gave rise to undifferentiated epiblast (*Ndrg2*, *Ndrg1*, *Cldn7*, *Cldn6, Cldn3*), surface ectoderm (*Tfap2a*, *Sfn*, *Krt18*, *Krt8*, *Cldn4*) or primordial germ cell (PGC) (*Sox17*, *Prdm14*, *Prdm1*, *Dnd1*, *Dppa3*) identities (Fig. 1E, F).

This analysis confirmed previous mesodermal and ectodermal annotations, demonstrated the formation of hematopoietic progenitors that a previous study had suggested (Morgani et al., 2018), and uncovered the presence of PGCs.

### t00-m2Dgas resemble the E5.5 epiblast

To better define the status of our colonies at t=0h, we compared their scRNAseq data to that of E5.5 embryos (Nowotschin et al., 2019). On the UMAP reduction plot, t00-Epi cells are localized close to E5.5-Epi cells, but separately from them (Fig. 2A), suggesting the presence of both similarities and differences. We first looked at the expression of 9 genes involved in pluripotency and Epi maturation (Fig. 2B). The only marked difference concerned *Dnmt3b*, whose expression was found to be lower in m2Dgas at 0h than in E5.5-Epi. DNMT3B is involved in *de novo* methylation during epiblast maturation but its absence can be partially compensated by DNMT3A. *Dnmt3b*^-/-^ embryos show no phenotype before E9.5 and *Dnmt3b*^+/-^ mice are viable and have no phenotype (Okano et al., 1999). Since *Dnmt3b* expression was captured in 98% of t00-Epi cells, its expression was probably sufficient to allow differentiation to proceed normally. To go further, we examined the expression of genes associated with the gene ontology (GO) term “*transcription regulator activity*” that are upregulated in the E5.5- Epi cluster compared to the two extra-embryonic clusters, E5.5-ExE and E5.5-VE (Fig. S2A). Similar to *Dnmt3b*, a few genes, such as *Kbm5b*, *Psip1*, *Tcf3* and Tead2, had a lower expression in t00-Epi cells compared to E5.5-Epi, despite being expressed by a similar fraction of these cell populations. We also performed unbiased GO analysis using the topgo pipeline (Alexa & Rahnenführer, 2009) on genes whose expression was upregulated in t00-Epi cells compared to E5.5-Epi. The most significant GO terms emerging from this analysis are related to metabolic processes (Fig. S2B). Among the upregulated genes (Fig. S2C) are members of the coiled-coil-helix-coiled-coil-helix domain (CHCHD) and mitochondrial ribosomal protein (MRP) families, both of which are involved in mitochondrial metabolism (Zhou et al., 2017; Huang et al., 2020). Overall, t00-Epi and E5.5-Epi have very similar gene expression profiles and the small differences we detected between them probably reflect metabolic differences between the *in vitro* culture system and the embryo.

**Figure 2.**
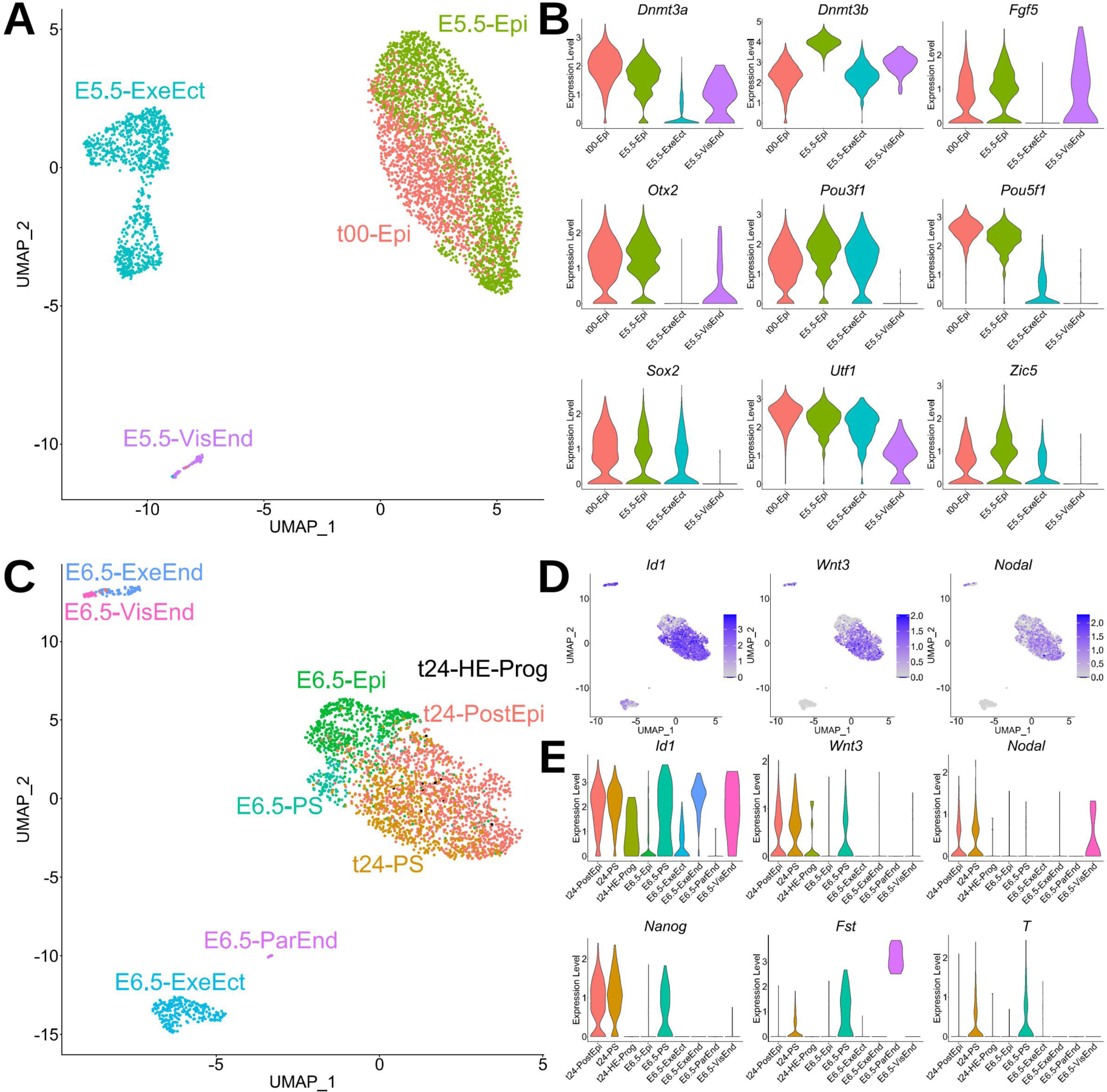
**Comparison of t=0h and t=24h BMP4-s;mulated m2Dgas cells with their embryonic counterparts at E5.5 and E6.5** ***(A)*** UMAP representa-on of the integrated dataset for t=0h m2Dgas cells and E5.5 mouse embryo cells (Nowotschin et al., 2019), showing a par-al overlap of t00-Epi and E5.5_Epi cells. ***(B)*** Violin plots of the expression of key genes involved in epiblast pluripotency. ***(C)*** UMAP representa-on of the integrated dataset for t24-m2Dgas cells and E6.5 mouse embryo cells (Pijuan- Sala et al., 2019), showing that m2Dgas cells are close but dis-nct from their embryonic counterparts. ***(D)*** Same UMAP plot showing the associated expression of Id1 (a BMP signaling target), Wnt3 and Nodal. ***(E)*** Violin plots of the expression of key genes in posterior epiblast. <u>Epi:</u> Epiblast, <u>PostEpi:</u> Posterior Epiblast, <u>UndEpi:</u> Undifferen-ated Epiblast, <u>PS:</u> Primi-ve Streak, <u>HE-Prog:</u> Haemato-Endothelial Progenitors, <u>ExeEct:</u> Extra-embryonic ectoderm, <u>ExeEnd:</u> Extra-embryonic Endoderm, <u>VisEnd:</u> Visceral endoderm.

### t24-m2Dgas are in a transition state between posterior epiblast and primitive streak

A similar comparison was carried out between t24-m2Dgas and E6.5 embryos (Pijuan-Sala et al., 2019). The t24-PostEpi cluster is close to E6.5-Epi, while t24-PS is close to E6.5-PS, but once again there is no perfect match between them (Fig. 2C). We first looked at signaling activities. *Id1* (a BMP target gene), *Wnt3*, and *Nodal* show fairly homogenous expression in the m2Dgas clusters (Fig. 2D), with slightly higher mean expression in t24-PS (Fig. 2E, FigS4). This is in agreement with data showing that after its initial homogenous activation in a differentiating m2Dgas, *Nodal* expression persists in the outer ring where the PS is formed, whereas it is downregulated in the rest of the colony (Plouhinec et al., 2022). *Nanog* is a direct target of ACTIVIN/NODAL signaling (L. T. Sun et al., 2014) and 24h after BMP exposure *Nodal*^-/-^ m2Dgas have lower *Nanog* expression than WT m2Dgas (Plouhinec et al., 2022). Consistent with this, *Nanog* is expressed in both m2Dgas clusters, but with a higher mean expression in t24-PS (Fig. 2E, Fig. S2D). Higher signaling activities, as well as higher *Nanog* expression (Turner et al., 2014), likely explain the induction of some PS markers (*Fst*, *T*) in t24-PS cells (Fig. 2D). However, other PS markers duly expressed in E6.5-PS, such as *Eomesodermin* (*Eomes)* and *Mixl1*, are not detected in t24-PS cells, suggesting that these cells may correspond to an early stage of PS development. At the same time, some genes associated with pluripotency start to be downregulated in t24-PS (*Otx2*, *Utf1*), while others are no longer expressed in both clusters (*Pou3f1*, *Sox2*). Interestingly, a few cells already show HE-Prog characteristics (Fig. 2C, Fig. S2E), which seems surprising given that no mesoderm has yet formed, but is consistent with the timing of their formation in the embryo, where they are detected as early as E6.75 (Pijuan-Sala et al., 2019)

### t48-m2Dgas recapitulate early steps in the formation of PGCs and posterior PS derivatives

To accurately identify the different cell types present on t48 m2Dgas (Fig. 3A), a cluster stability analysis (Fig. S3A) was carried out. A resolution of 1.1 was chosen because it allowed the identification of an ExeMes cluster, associated with a specific upregulation of *Foxf1*, *Hand1* and *Bmp4,* and of a PGC cluster, associated with a specific upregulation of *Prdm1* (*Blimp1*), *Prdm14* and *Dnd1* (Fig. S3B), all of which are involved in the formation of PGCs (Ohinata et al., 2005; Kurimoto et al., 2008; Li et al., 2018).

**Figure 3.**
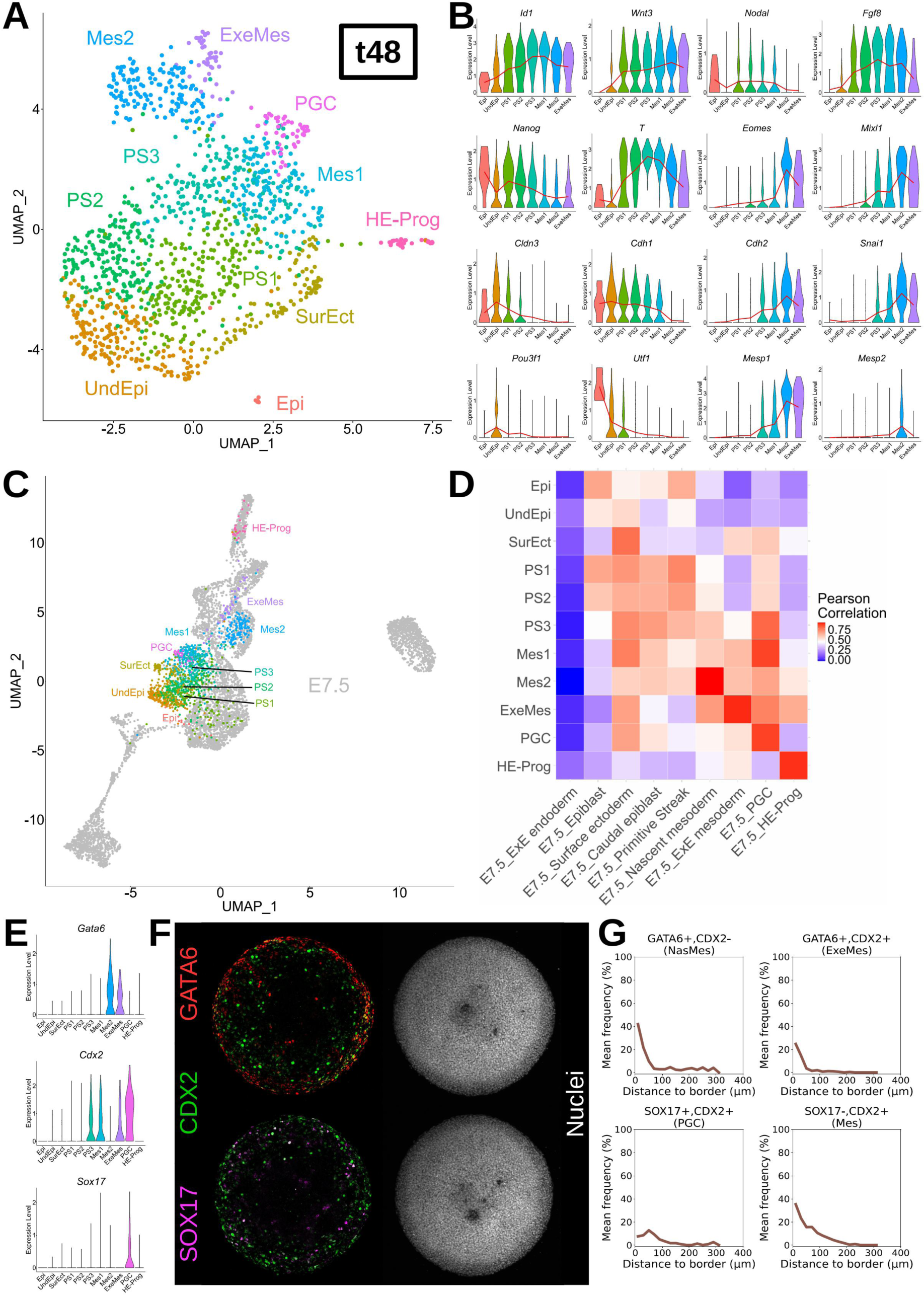
**Comparison of t=48h BMP4-s;mulated m2Dgas cells with their embryonic counterparts at E7.5** ***(A)*** UMAP representation of the different cell identity clusters annotated in BMP-stimulated m2Dgas at t=48h. ***(B)*** Violin plots of the expression of key genes involved in primi-ve streak forma-on, EMT and mesodermal differen-a-on. The red line tracks the mean expression of the gene in each cluster. ***(C)*** UMAP representation of the integrated dataset for t48-m2Dgas cells and E7.5 mouse embryo cells (Pijuan- Sala et al., 2019). Only the m2Dgas clusters are colored and labelled. See Fig. S3C for the annotation of the embryonic clusters. ***(D)*** Mean Pearson correlation in the batch-corrected PCA space between m2Dgas and embryonic clusters. ***(E)*** Violin plots of the expression of Gata6, Cdx2 and Sox17 in t48-m2Dgas clusters. Gata6 and Cdx2 are co- expressed in ExeMes. Cdx2 and Sox17 are co-expressed in PGCs. ***(F)*** Maximum intensity projections of a t48-m2Dgas (700 µm) immuno-stained for GATA6, CDX2 and SOX17. Single cell co-expression analysis of m2Dgas shown in **(F)**. Frequencies are averaged along the colony radii (n=3 for each curve).

As the main clusters present at t=48h are Epi, PS and Mes, we first focused on the events leading to the formation of mesoderm and looked at key gastrulation genes (Fig. 3B). A notable aspect of PS formation is that it is the point of convergence of several signaling pathways (Morgani & Hadjantonakis, 2020). Here, from UndEpi to PS3, the expression of *Id1*, *Wnt3*, *Nodal* and *Fgf8* shows a gradual activation, but this expression decreases in ExeMes.

The correct specification of the PS is also dependent on several transcription factors, including NANOG, T, EOMES and MIXL1 (Tosic et al., 2019; Turner et al., 2014; Ng et al., 2005). Like in t24-m2Dgas, *Nanog* expression appears to precede that of *T*, because it peaks in PS1 whereas *T* expression peaks in PS3. In contrast, *Eomes* and *Mixl1* are gradually activated in the PS clusters, and only reach peak expression in the Mes2 cluster. These changes coincide with an epithelial–mesenchymal transition (EMT) that involves a loss of tight junctions and a SNAI1- dependent switch from E-CADHERIN (*Cdh1*) to N-CADHERIN (*Cdh2*) (Reviewed in Nakaya and Sheng, 2008; Bardot and Hadjantonakis, 2020). These events take place in the PS of m2Dgas, as *Cldn3* and *Cdh1* are downregulated as mesoderm forms, while conversely, *Cdh2* and *Snai1* are upregulated. During this transition, expression of the pluripotency factors *Pou3f1* and *Utf1* is also lost, while expression of the mesodermal markers *Mesp1* and *Mesp2* is activated in Mes clusters.

To better characterize the annotated clusters, we compared t48-m2Dgas to the E7.5 embryo (Pijuan-Sala et al., 2019) (Fig. 3C, Fig. S3C). This time, as the number of cell types is higher, the mean correlation coefficient was computed in the batch-corrected PCA space between m2Dgas and embryonic cell types (Fig. 3D). The m2Dgas clusters SurEct, Mes2, ExeMes, PGC and HE-Prog have a high correlation to their embryonic counterparts, the E7.5 Surface ectoderm, Nascent mesoderm, ExE mesoderm, PGCs and haemato-endothelial progenitors (HE-Prog), respectively, suggesting that they have similar gene expression profiles. This is also consistent with their overlapping locations on the UMAP (Fig. 3C, Fig. S3C). In contrast, the Epi and PS clusters do not merge with E7.5 Epiblast and Primitive Streak on the UMAP, and have lower correlation coefficients. Interestingly, these discrepancies between the Epi and the PS of m2Dgas and their *in vivo* counterparts, do not prevent them from giving rise to derivatives that are much closer to their *in vivo* counterparts.

Finally, we determined where on the m2Dgas some of the annotated clusters were located. To locate the PS, we performed *in situ* hybridizations (ISH) for *Nodal* and *T*, which are both expressed there (Fig. 3B). As before (Plouhinec et al., 2022), *Nodal* expression was found to be restricted to an inner ring, its inner-most boundary coinciding with that of *T* (Fig. S3D). We then searched for specific combinations of markers to identify Mes2, ExeMes and PGCs. The Mes2 cluster can be identified as *Gata6+Cdx2-*, the ExeMes cluster as *Gata6+Cdx2+*, and the PGCs as *Sox17+Cdx2+* (Fig. 3E). IF were thus performed using these combinations (Fig. 3F), and the resulting images were quantified at the level of single nuclei (Fig. 3G). Mes2, ExeMes and PGCs were located within a ring bordering the colonies, in agreement with the expression of *Bmp4*, as detected by ISH (Fig. S3D), and also with the fact that this is where PS formation is known to begin (Morgani et al., 2018). To confirm the presence of PGCs in m2Dgas, we checked whether some of these cells also expressed SOX2. In the embryo, PGCs reactivate SOX2 expression shortly after their formation (Yamaji et al., 2008), and it is critical for their maintenance (Campolo et al., 2013). SOX2 expression was indeed detected in a ring of cells at the border of t60-m2Dgas (Fig. S3E), and most SOX2+ nuclei in this region were also CDX2+, attesting of their PGC identity (Fig. S3F).

### t72-m2Dgas show PGCs maintenance and posterior mesodermal diversification

When characterizing the cell identities present on t72-m2Dgas (Fig. 4A), we found that setting the resolution at 1.3 allowed the identification of a PGC cluster (Fig. S4A), associated with a specific expression of *Prdm1* (*Blimp1*)(Fig. 4B). This analysis led to the identification of 2 Epi, 1 SurEct, 6 mesoderm (including ExeMes, Mesenchyme and Allantois), 1 PGC and 1 HE-Prog clusters.

**Figure 4.**
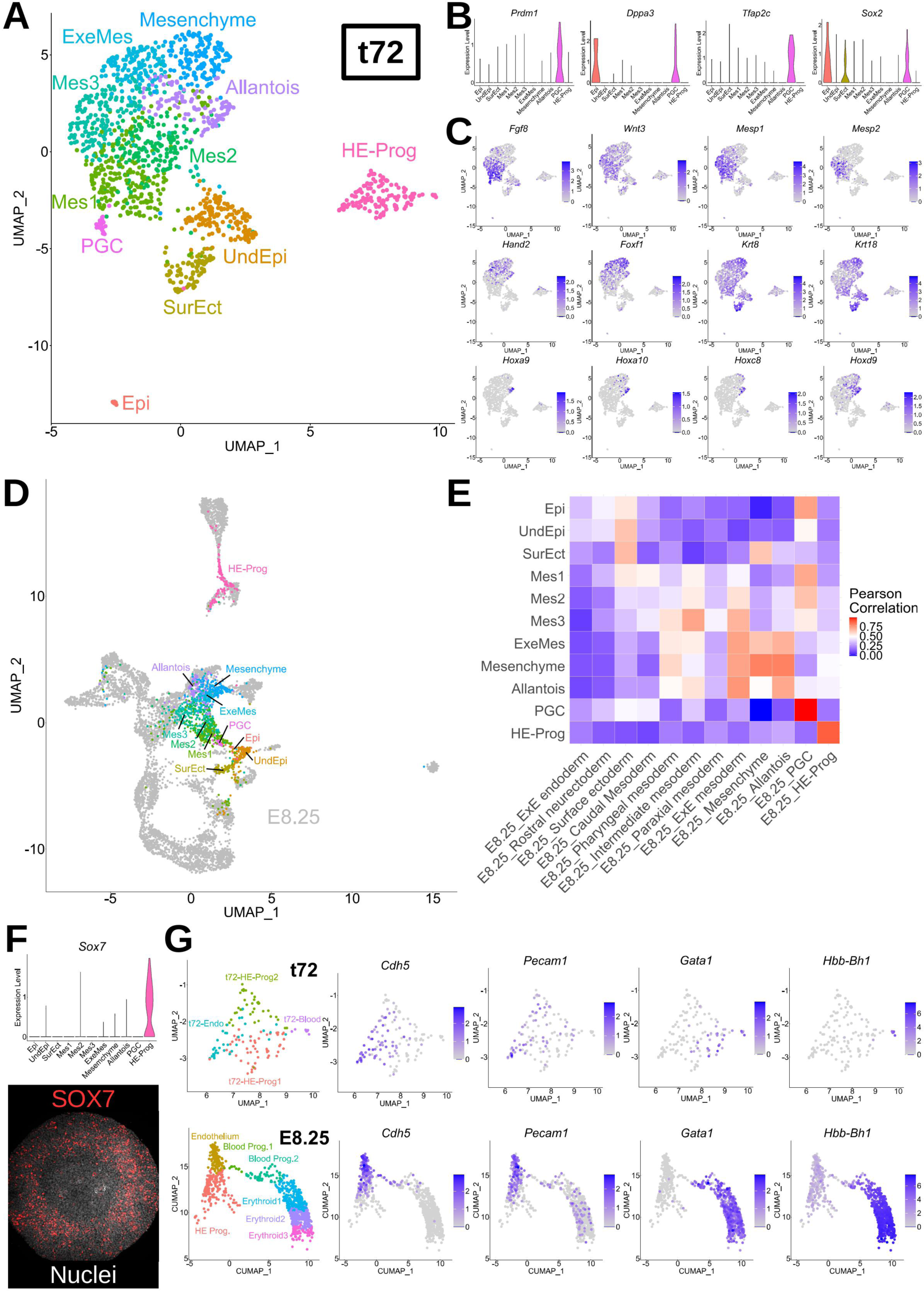
**Comparison of t=72h BMP4-s;mulated m2Dgas cells with their embryonic counterparts at E8.25** ***(A)*** UMAP representa-on of the cell iden-ty clusters annotated in BMP4-s-mulated m2Dgas at t=72h. ***(B)*** Violin plots of the expression of key genes involved in the maintenance of PGCs. ***(C)*** Same UMAP than in (A) showing the associated expression of genes involved in mesoderm forma-on. ***(D)*** UMAP representa-on of the integrated dataset for t72-m2Dgas cells and E8.25 mouse embryo cells (Pijuan- Sala et al., 2019). Only the m2Dgas clusters are colored and labelled. See Fig. S4C for the annota-on of the embryonic clusters. ***(E)*** Mean Pearson correla-on in the batch-corrected PCA space between m2Dgas and embryonic clusters. ***(F)*** (top) Violin plot of Sox7 expression. (bo9om) Maximum intensity projec-ons of a t72-m2Dgas (700 µm) immunostained for SOX7. UMAP representa-ons of Haemato-Endothelial progenitors from t72-m2Dgas and E8.25 embryos and the associated expression of genes involved in endothelial (Cdh5, Pecam1) or blood (Gata1, Hbb-Bh1) commitment.

Once formed, PGCs depend not only on *Sox2* for maintenance, but also on *Dppa3* (*Stella*) and *Tfap2c* (*Ap2γ*) (Nakashima et al., 2013; Weber et al., 2010; Campolo et al., 2013). All three gene are co-expressed in t72-m2Dgas PGCs (Fig. 4B), showing that those that formed earlier have progressed further.

We then focused on the characteristics of the mesodermal clusters. The correct early mesodermal specification relies on FGF and WNT signaling (Arkell et al., 2013; X. Sun et al., 1999), and requires also *Mesp1* and *Mesp2* (Ajima et al., 2021)*. Fgf8*, *Wnt3*, *Mesp1* and *Mesp2* are expressed in the Mes1-3 clusters (Fig. 4C), which show features of nascent mesoderm (Pijuan-Sala et al., 2019). From there, the production of extra-embryonic mesoderm and its derivatives is dependent on *Hand2* (Yamagishi et al., 2000), *Foxf1* (Mahlapuu et al., 2001), *Hox* genes (Scotti & Kmita, 2012) and KERATIN filaments (Nahaboo et al., 2022). In the mouse gastrulation atlas (Pijuan-Sala et al., 2019), *Hand2* and *Foxf1* are expressed in extra-embryonic mesoderm, mesenchyme and allantois. *Krt* genes are upregulated in the mesenchyme, while some *Hox* genes are expressed in the allantois. All these specific gene regulations were also observed in m2Dgas (Fig. 4C).

To position the annotated clusters relative to their *in vivo* counterparts, we compared t72- m2Dgas to E8.25 embryo (Pijuan-Sala et al., 2019) (Fig. 4D, Fig. S4C), and calculated their mean correlation coefficients in a batch-corrected PCA space (Fig. 4E). On the UMAP, the m2Dgas SurEct, ExeMes, Mesenchyme, Allantois, PGCs and HE-Prog clusters are globally close to at least a subset of cells from their embryonic counterpart, whereas this is not the case for the Mes1-3 clusters. This is also reflected in their correlation score (Fig. 4E). To confirm these results, we examined the expression of key genes involved in the specification of embryonic mesoderm. These included *Nkx1.2*, as a marker of caudal epiblast/mesoderm (Rodrigo Albors et al., 2018), *Nkx2.*5 for cardiac mesoderm (Lints et al., 1993), *Osr1* for intermediate mesoderm (Mugford et al., 2008), *Foxc2* for paraxial/somatic mesoderm (Wilm et al., 2004), as well as other marker genes picked from (Pijuan-Sala et al., 2019). In contrast to the embryo, none of them were expressed in t72-m2Dgas (Fig. S4D), confirming that no embryonic mesoderm is properly established in BMP4-stimulated m2Dgas at this time point.

Finally, we focused on the haemato-endothelial progenitors (HE-Prog) that are present in t72- m2Dgas. *Sox7*, which has been associated with the onset of vasculogenesis and angiogenesis (Lilly et al., 2017), was found to be specifically expressed in cells from the HE-Prog cluster (Fig. 4F). Its expression pattern in m2Dgas was therefore investigated. SOX7-positive cells were found within a wide ring of tissue beginning at the edge of the colonies (Fig. 4F). HE-Progs give rise to either endothelial or blood cells. *Cdh5* (*VE-Cadherin*) and *Pecam1* (*Cd31*) are important for the function of endothelial cells (Carmeliet et al., 1999; Pinter et al., 1997), while the transcription factor *Gata1* is required for the maturation of blood cells and the expression of Haemoglobin genes (Pevny et al., 1991; Barile et al., 2021). We looked at the expression of these genes in the HE-Prog cluster and compared it to their expression in the embryo (Fig. 4G). In a manner reminiscent of what happens in the embryo, both *Cdh5* and *Pecam1* were expressed in a sub-cluster of cells that we named t72-Endo, while *Gata1* and *Hbb-Bh1* were expressed in another sub-cluster that we named t72-Blood. Examination of the correlation coefficients between these m2Dgas cells and their putative embryonic counterparts, and a comparison of the most upregulated transcription factors in these cell populations validated these annotations (Fig. S4E-F).

### Reducing BMP4 exposure enables the formation of anterior Primitive Streak

Our transcriptome analyses of BMP4-stimulated m2Dgas confirmed that anterior PS and its derivatives are absent, as described in previous studies (Morgani et al., 2018; Plouhinec et al., 2022), and also demonstrated that the cellular identities produced are restricted to those forming in the most proximal region of the gastrulating embryo. BMP4 concentration was maintained at 50ng/ml for 48h in these experiments. We wondered whether reducing BMP4 concentration or exposure time would allow the production of more distal cell identities. Treatments of mESCs with concentrations as low as 1ng/ml of BMP4 have been reported (Najafi et al., 2009; Makoulati et al., 2009), and 6h stimulation of m2Dgas with 50 ng/ml of BMP4 was found sufficient to activate *Wnt3* expression, which can then promote its own expression (Plouhinec et al., 2022). m2Dgas were therefore exposed to 3 conditions: 1ng/ml of BMP4 for 48h (B1-48h); 50ng/ml for 6h (B50-06h); 50ng/ml for 48h (B50-48h). To provide a control for anterior PS formation, m2Dgas were exposed to WNT3A for 48h (W-48h) (Fig. 5A). We first checked BRA expression at 24h. The B50-06h and B50-48h conditions led to similar patterns of expression, whereas the B1-48h condition resulted in lower BRA expression (Fig. S5A). The expression of the *Bra* gene was similarly affected by these conditions, even though they had no detectable impact on that of *Wnt3* and *Nodal* (Fig. S5B). As a key feature of anterior PS is to be triple positive for BRA, FOXA2 and GSC (Morgani et al., 2018; Pijuan-Sala et al., 2019), we examined the expression of these factors at t=48h (Fig. 5B). Conditions B1- 48h and B50-06h resulted in similar expression patterns, with, compared to condition B50- 48h, no change for BRA, a larger inner ring of FOXA2 expression, and a global increase of GSC expression. To better identify and position the cell identities present on these m2Dgas, we measured the expression of each marker in individual nuclei. Posterior PS was defined as FOXA2-/BRA+/GSC+, anterior PS as FOXA2+/BRA+/GSC+, and epiblast as BRA-/GSC- (Fig. 5C).

**Figure 5.**
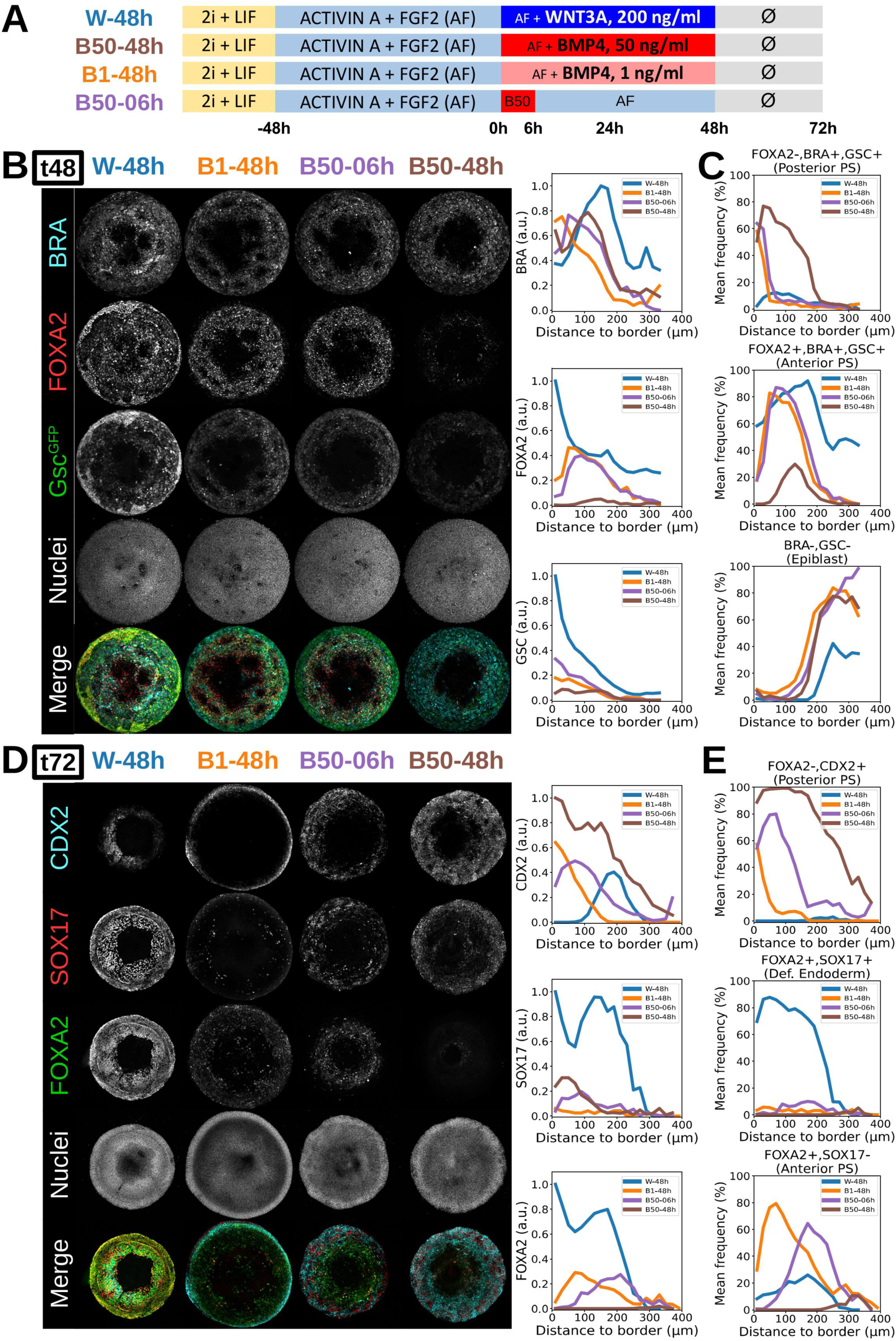
Reducing m2Dgas exposure to BMP4 allows them to form both posterior and anterior PS deriva;ves. **(A)** Timeline of the culture conditions used to test the effect of a reduction in BMP4 exposure on m2Dgas development. ***(B)*** Maximum intensity projections of m2Dgas (700 mm) treated as defined in (A) and immunostained aQer 48h of differen-a-on. Adjacent graphs show fluorescence intensity levels normalized by Hoechst and averaged along the colony radii (n=4 for each curve). ***(C)*** Single cell co-expression analysis of m2Dgas shown in (B). Frequencies are averaged along the colony radii (n=4 for each curve). ***(D)*** Maximum intensity projections of m2Dgas (700 mm) treated as above and immuno-stained aQer 72h of differen-a-on. Adjacent graphs show fluorescence intensity levels normalized by Hoechst and averaged along the colony radii (n=4 for each curve). Single cell co-expression analysis of m2Dgas shown in **(D)**. Frequencies are averaged along the colony radii (mean ± SEM, n=4).

A posterior PS is still formed at the edge of B1-48h and B50-06h colonies, but the ring is thinner than that in B50-48h colonies. The presence of pSMAD1 at the edge of the colonies (Fig. S5C) and the induction of *Bmp4* and *Hand1* expression (Fig. S5D), attest to the BMP signaling activity underlying posterior mesoderm formation. The inner ring of FOXA2 expression observed in B1-48h and B50-06h colonies was found to contain predominantly anterior PS cells (Fig. 5B, C), a few of which expressed the key DE marker SOX17 (Burtscher & Lickert, 2009) (Fig. S5E-F). In B50-48h m2Dgas, FOXA2 is expressed at t=48h but its expression is not maintained at t=72h (Morgani et al., 2018; Plouhinec et al., 2022). In contrast, we found that both B1-48h and B50-06h colonies still expressed FOXA2 at t=72h, and some DE cells were still present (Fig. 5C-D). In addition, CDX2+/FOXA2- nuclei were present at the edge of these colonies, indicating that they had also formed posterior PS derivatives.

Reducing the exposure of m2Dgas to BMP4 is therefore sufficient to allow them to form and to maintain a PS that produces both posterior and anterior cellular identities.

### Timed inhibition of BMP signaling allows the formation of paraxial mesoderm

Since BMP signaling was found to be active in B1-48h and B50-06h m2Dgas well beyond their window of exposure to recombinant BMP4, we wanted to assess the impact on m2Dgas development of restricting the activity of the pathway to the first 24h. To do this, we treated the colonies with a pharmacological inhibitor of BMP signaling, LDN193189, hereafter designated Bi, from t=24h until the end of the culture (Fig. 6A). We characterized the impact of the treatment at t=72h, first looking at pSMAD1 to check that BMP signaling had indeed been abolished (Fig. S6A). Although inhibition of BMP signaling resulted in the elimination of pSMAD1 from the entire surface of the colonies, expression of the neurectoderm marker SOX1 remained restricted to their center, indicating that the limited exposure of the surrounding cells to BMP4 was sufficient to commit them to a different fate (Fig. S6B).

**Figure 6.**
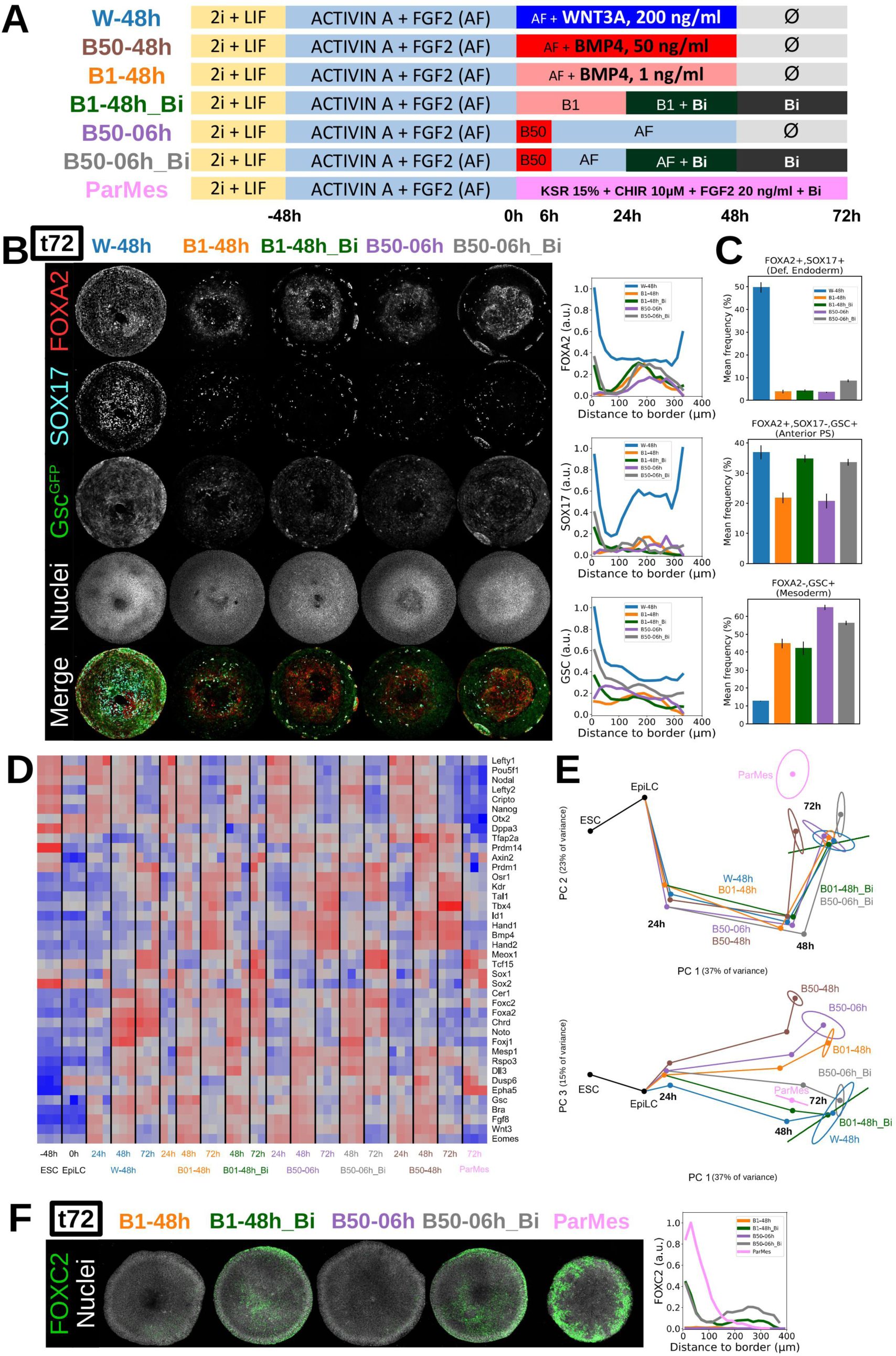
**: Inhibi;on of BMP signaling 24h aVer its induc;on in m2Dgas leads to the forma;on of paraxial mesoderm** ***(A)*** Timeline of the culture conditions used to test the impact of BMP signaling inhibi-on on m2Dgas. ***(B)*** Maximum intensity projections of m2Dgas (700 mm) treated as defined in (A) and immunostained aQer 72h of differen-a-on. Adjacent graphs show fluorescence intensity levels normalized by Hoechst and averaged along the colony radii (n=4 for each curve). ***(C)*** Single cell co-expression analysis of m2Dgas shown in (B). Frequencies are averaged along the colony radii (n=4 for each curve). ***(D)*** Gene expression matrix obtained via RT-qPCR of pooled colonies at similar -me points for each of the treatments shown in (A). ***(E)*** Projection of the RT-qPCR data shown in (D) in the space defined by the first two principal components of the dataset (top), or by its first and third components (bo9om), showing the developmental trajectories followed by m2Dgas for each treatment. Maximum intensity projec-ons of m2Dgas (700 mm) treated as above, and immuno-stained for FOXC2 aQer 72h of differen-a-on. Fluorescence intensity levels were normalized to Hoechst and averaged along the colony radii (n=4 for each curve). BMP inhibition results in the expression of FOXC2, a paraxial mesoderm marker, in the outer ring of the colonies, similar to ParMes-inducing condi-ons.

**Figure 7:**
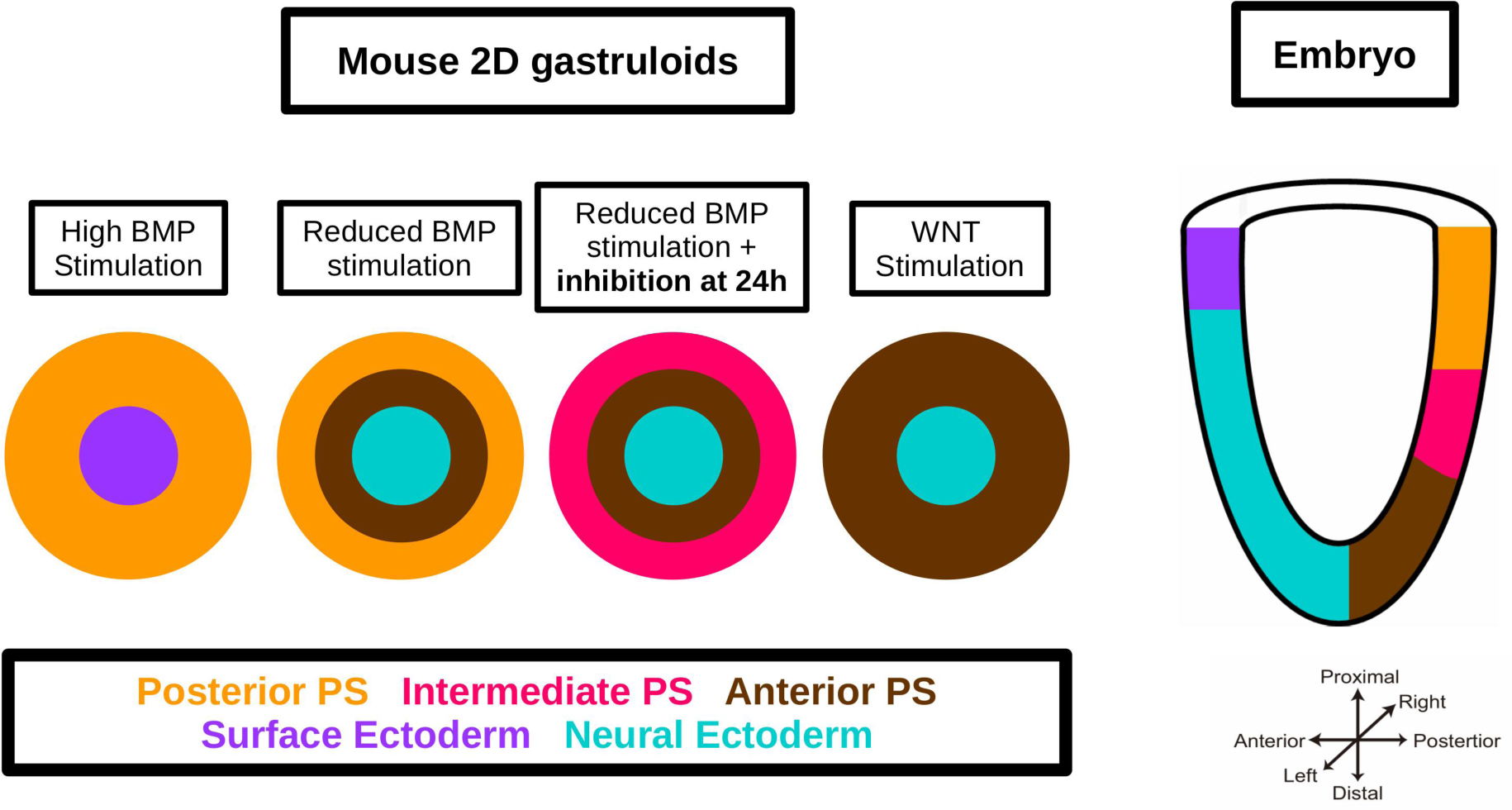
Graphical summary High and con-nuous BMP s-mula-on of m2Dgas leads to the forma-on of iden--es corresponding to the proximal part of the mouse embryo: surface ectoderm in the center, and posterior PS deriva-ves at the border. In contrast, WNT s-mula-on leads to the forma-on of distal iden--es: neural ectoderm in the center, and anterior PS deriva-ves at the border. Reduced exposure to exogenous BMP, either in -me or concentra-on, allows the forma-on and maintenance of an inner ring of anterior PS while maintaining an outer ring of posterior PS. Reduced BMP exposure also results in neural ectoderm forming in the center of the colonies instead of surface ectoderm. Inhibi-on of BMP signaling, aQer an ini-al 24h exposure to BMP4, does not lead to a spa-al organiza-on similar to the WNT s-mula-on. Instead, deriva-ves of the intermediate part of the PS, such as paraxial mesoderm, form at the colony border. Embryo schema-c adapted from (Peng et al., 2016). PS: primi-ve streak.

We then looked at the formation of anterior PS (aPS) and DE. Inhibition of BMP signaling resulted in the presence of small patches of FOXA2 expression along the edge of the colonies, forming an incomplete outer ring (Fig. 6B). These patches were also positive for SOX17 and GSC, indicating that they contained DE cells, but overall, the inhibition had little impact on the total number of DE cells (Fig. 6C). Interestingly, the parts of the outer ring that remained negative for FOXA2 still expressed GSC, as was the case in uninhibited colonies, suggesting that mesoderm had formed there (Fig. 6 B, C). *Gata6* is known to be expressed in aPS, NasMes, ExeMes and lateral mesoderm. The same is true for *Krt8*, except that it is not expressed in NasMes (Pijuan-Sala et al., 2019). Upon BMP4 inhibition, GATA6 remained expressed in the outer part of the colonies, while KRT8 was lost, except in the small patches of DE (Fig. S6E), suggesting the presence of something still resembling mesoderm.

To go further, we performed RT-qPCR of whole colonies collected at different time points, looking at a greater number of markers (Fig. 6D). We also adapted a protocol from previous studies (Gouti et al., 2014; Chal et al., 2015), and used it as a condition to induce paraxial mesoderm (ParMes; Fig. 6A). Principal Component Analysis (PCA) of the resulting gene expression matrix (Fig. 6D) was used to compare the developmental trajectories obtained for each condition (Fig. 6E). The PC1-PC2 plot mainly separates ParMes from all the other expression profiles obtained at 72h, which appear close to each other. In contrast, the PC1- PC3 plot distributes these different expression profiles along an arc, with B50-48h at one end, B1-48h and B50-6h close together at an intermediate position, and B1-48h_Bi and B50-6h_Bi grouped with W-48h at the other end. ParMes is closer to the latter group, but at a distance. We then took a closer look at the expression of certain genes (Fig. S6D). The mesodermal genes *Mesp1* and *Kdr* remained expressed at 48h after BMP signaling inhibition, while *Foxc2*, a key regulator of paraxial mesoderm specification (Wilm et al., 2004), was upregulated. These 3 genes were weakly expressed 48h after the start of WNT stimulation, suggesting that the initial 24h exposure to BMP4 was responsible for these expressions. Expression of *Meox1*, another key regulator of paraxial mesoderm (Candia et al., 1992; Mankoo et al., 2003), was then activated at t=72h, following that of *Foxc2*. To locate these cells on m2Dgas, we examined the expression pattern of FOXC2 (Fig. 6F). FOXC2 was only detected after inhibition of BMP signaling and was found to be expressed mainly at the edge of the colonies, confirming that mesoderm had formed there, but lower expression levels were also observed in the inner region of the colonies.

BMP signaling inhibition also led to the upregulation of genes involved in axial mesoderm/ notochord formation, such as *Noto* and *Foxj1* (Ben Abdelkhalek et al., 2004; Alten et al., 2012), in a dynamic similar to that induced by WNT3A stimulation, while expression of extra- embryonic mesoderm markers (*Hand1*, *Hand2* and *Bmp4*) was lost. The key regulator of HE- Prog formation, *Tal1,* remained expressed, but the loss of SOX7 expression implied that the specification of HE-Prog was defective (Fig. S6E).

We conclude that cells exposed to BMP signaling at the onset of PS formation are primed for mesoderm formation. Once the PS has formed, the maintenance of BMP signaling is essential for the formation of ExeMes and HE-Prog, but its elimination allows the emergence of more distal mesoderms.

## Discussion

Our results show that exposure to BMP4 is the most critical input for PS patterning, as its modulation determines which combination of cell identities emerges from the PS. These results are consistent with what is known already of the conditions associated with the formation of specific identities, such as PGCs or paraxial mesoderm, in the mouse embryo. This further supports the idea that, despite their limited physical similarity to the post-implantation embryo, m2Dgas are an excellent model for studying cell fate specification during gastrulation.

Our scRNA-seq analysis of m2Dgas exposed to high levels of BMP4 for up to 48h reveals that they form all the cell types that normally derive from the proximal-most region of the epiblast.

This region is continuously exposed to the BMP4 and BMP8b ligands produced first by the adjacent EXE, and then by the extra-embryonic mesoderm to which it gives rise. Interestingly, it is after 48h of exposure to BMP4 that the cell identities present on m2Dgas appear to be closest to their embryonic counterparts. This is somewhat reminiscent of the similarities observed between gastrulating embryos of different vertebrate species, which led to the concept of an hourglass-like bottleneck for vertebrate development at this stage (Irie, 2017; Mayshar et al., 2023). While the later divergence in m2Dgas may reflect the limitations of the system to recapitulate patterning events beyond a certain time in culture, the early convergence suggests that the distinct cell identities that gradually emerge on these colonies are, like their embryonic counterparts, interdependent for their development, and therefore face greater constraints on their gene expression than the EpiLC progenitors from which they are derived.

The case of PGCs may illustrate this point. The PGCs we found on BMP4-induced m2Dgas 48h and 72h after induction are very close to those present in E7.5 and E8.25 embryos. Embryo studies have shown that BMP signals, which come in the form of BMP4 and BMP8b from the ExE, and BMP2 from the VE, are required to induce PGC formation (Lawson et al., 1999; Ying et al., 2000; Ying & Zhao, 2001). However, recent studies have shown that as PGCs emerge in the budding allantois of the embryo, they become refractory to BMP signaling, and have lower levels of pSMAD1 than neighboring cells (Senft et al., 2019; Morgani & Hadjantonakis, 2021). One of these studies also points to the possible involvement of a non-canonical mode of action for BMP with respect to PGC specification (Morgani & Hadjantonakis, 2021). Since *Smad1* and *Smad5* mutants form reduced numbers of PGCs (Arnold et al., 2006; Chang et al., 1999; Tremblay et al., 2001), all of this may suggest that BMP signaling is required in cells playing a critical role in the emergence of PGCs as part of a niche, but not in PGCs themselves. The production of PGCs *in vitro* currently involves treating EpiLCs with BMP4 and BMP8b, as well as LIF, SCF and EGF (Hayashi et al., 2011). Their production in m2Dgas requires only prolonged exposure to BMP4, which appears sufficient to cover all BMP requirements identified in the embryo (Ying et al., 2000). The analysis of available scRNA-seq datasets failed to detect significant expression levels of LIF, SCF and EGF in the cells that are surrounding pre-PGCs and PGCs in the PS and in the extra-embryonic mesoderm, both in the embryo and in m2Dgas (Pijuan-Sala et al., 2019; this study). Key aspects of the emergence of PGCs, and the way in which cells surrounding them contribute to this emergence and support their development, have yet to be elucidated. Our results suggest that m2Dgas provide a convenient model system to further dissect the interactions that lead to the formation of PGCs.

After 48 hours of exposure to BMP4, m2Dgas also gave rise to distinct endothelium and blood progenitors, as the posterior primitive streak does in the embryo. The origin of these progenitors, and the extent to which they share common ancestors, has been the focus of countless studies, both *in vitro* and *in vivo* (Vink et al., 2022). One of the most recent, relying on single cell lineage tracking by *in vivo* barcoding to resolve their cellular genealogy (Biben et al., 2023), actually showed that these lineages arise from a trio of precursors: the haemangioblast, already known from previous studies, which produces haematopoietic lineages and endothelium, the mesenchymoangioblast, which produces mesenchyme and endothelium, and the haematomesoblast, a novel haematogenic precursor that produces haematopoietic lineages and mesenchyme. There is evidence that these precursors emerge in the posterior PS quite rapidly after the onset of gastrulation. Although we were able to detect the presence of haemato-endothelial progenitors as early as 24h after the start of BMP4 exposure, the data we obtained did not allow us to identify separate identities among these progenitors before the emergence of distinct haemato and endothelium lineages alongside mesenchyme. The presence of these lineages on our colonies confirms that key cell fate specification mechanisms operating in the posterior PS are recapitulated in m2Dgas exposed to BMP4 for 48h.

However, other mesodermal derivatives that normally arise from BMP-exposed proximal PS territories to contribute to head and heart development fail to form properly in m2Dgas exposed to BMP4 for 48h. Although the Mes2 cluster identified at t48 appears to be a good match for the nascent mesoderm identified in the E7.5 embryo, none of the clusters identified at t72 match the caudal mesoderm or pharyngeal mesoderm cell populations identified at E8.25 (Pijuan-Sala et al., 2019). This is in marked contrast to the PGC and HE progenitor populations, which at this stage are still close to their *in vivo* counterparts. HE progenitors and PGCs have in common that they are among the earliest cell identities to differentiate in the proximal PS and in BMP4-treated m2Dgas. Their persistence in such m2Dgas at t72 suggests that the conditions within these colonies remain supportive of their developmental progression, but not that of other proximal PS derivatives. *Mesp1* plays a key role in initiating the program that leads to the formation of the caudal mesoderm and pharyngeal mesoderm identities, but its expression in their precursors is normally transient, and *Mesp1* has been shown to first stimulate and then inhibit its own expression (Bondue et al., 2008). *Mesp1* is strongly expressed in Mes2 at t48, but it is still expressed there at t72, and may therefore disrupt the developmental progression of this cell population. *In vitro*, exposure of mESCs to high levels of BMP4 has been shown to promote the formation of HE progenitors, but also to prevent the formation of cardiomyocytes (Kattman et al., 2011). This suggests that the continuous exposure of Mes2 cells to BMP4, whose action, unlike in the embryo, is not locally counteracted by the production of BMP antagonists such as CHORDIN and CER1 by neighboring cells, may be a possible reason for the failure of such mesodermal derivatives to emerge on our colonies.

Modulating the activity of the BMP signaling pathway did allow the emergence of a wider range of cell identities on BMP4-treated m2Dgas. Reducing the concentration of the morphogen or the duration of exposure of the colonies to it resulted in the formation and maintenance of an inner ring of aPS containing even a few DE cells, while still maintaining an outer ring of cells with a posterior PS identity. However, it was only when an inhibitor of BMP signaling was added at t24 that a broad increase in the expression of paraxial and axial mesoderm markers was detected. The formation of cells with a paraxial mesoderm identity in these conditions is again consistent with an *in vitro* protocol that relies on both the activation of canonical WNT signaling and the inhibition of BMP signaling to differentiate mESCs into presomitic mesoderm cells (Chal et al., 2018). Inhibition of BMP signaling in m2Dgas presumably provides the relief from BMP influence that some PS cells get in the embryo when they are sufficiently removed from the proximal source of BMP and/or exposed to the BMP-signaling attenuating influence of the adjacent VE (Yang et al., 2010). Consistent with this, the inhibition of BMP signaling also led to the activation of *Chordin* and *Cer1* expression, which in the embryo begins in the aPS at mid-streak stage to ensure the emergence of cell identities otherwise blocked by BMP signaling (McMahon et al., 1998; Biben et al., 1998).

*Chordin* and *Cer1* are known NODAL targets (Reid et al., 2012), and their activation may also depend on the presence of other signals, such as WNT and FGF, which can also promote NODAL signaling and/or reduce BMP signaling (Ben-Haim et al., 2006; Aubin et al., 2004; Sapkota et al., 2006). In m2Dgas, BMP signaling promotes the expression of *Wnt3* and *Nodal*, but the activation of *Chordin* and *Cer1* requires its subsequent inhibition, highlighting its repressive effect on the expression of these NODAL and WNT targets. In our previous m2Dgas study, we found that BMP signaling has a differential effect on NODAL signaling targets, preventing the expression of some while leaving others unaffected or even promoting their expression (Plouhinec et al., 2022). It has been shown in other contexts that when both signaling pathways compete for limited amounts of a common component, such as SMAD4, SMAD1/5/9-dependent BMP signaling can reduce and limit the output of SMAD2/3- dependent ACTIVIN/NODAL signaling (Furtado et al., 2008; Pereira et al., 2012; Yamamoto et al., 2009). However, some SMAD2/3 targets that do not require SMAD4 for their expression have been identified (David & Massagué, 2018), and a recent study suggests that knocking out SMAD4 in mESCs prevents them from expressing BMP signaling targets, but has a lesser effect on the expression of NODAL signaling targets and even leaves some of them, such as *Lefty2*, unaffected (Guglielmi et al., 2021). This suggests that some NODAL signaling targets are sensitive to potential competition with BMP signaling, while others whose expression is independent of SMAD4 are not. This could explain, at least in part, the different effects of BMP exposure on the NODAL targets we included in our panel, from *Lefty2*, strongly activated, to *Chordin*, repressed, and finally the m2Dgas differentiation pattern we obtained for each treatment.

A recent study showed that the T-box transcription factors BRA and EOMES, whose induction in the posterior Epi by WNT and NODAL signaling, respectively, marks the onset of gastrulation, act at the level of chromatin to repress pluripotency and neurectoderm- promoting enhancers and to open mesoderm and endoderm-promoting ones (Tosic et al., 2019). Interestingly, BRA and EOMES are dispensable for the formation of pre-PGCs, but not for the formation of the PS-derived niche required for their maturation (Senft et al., 2019). The enhancers that depend on BRA and EOMES to become accessible are likely to include those that are going to mediate the influence of BMP and NODAL signaling on PS patterning. Although much of the action of the two pathways will involve their interaction with their transcription factor partners at these enhancers, an analysis of the SMAD2/3 interactome in human ESCs revealed that, in addition to their role in transcription, SMAD2/3 appear to be involved in many other cellular processes through their interaction with a number of protein complexes (Bertero et al., 2015). Although, to our knowledge, a similar study has not yet been performed for SMAD1/5/9, these results raise the question of whether BMP signaling may also affect these other aspects of NODAL signaling action.

The study of m2Dgas has provided welcome insights into the respective roles of the NODAL, WNT and BMP signaling pathways in patterning the primitive streak. We expect that the use of this *in vitro* model, as well as that of 3D gastruloids, will continue to advance our understanding of the mechanisms underlying the specification and differentiation of PS derivatives. Further progress will require a more detailed examination of the effects of graded increases in BMP exposure on cell fate specification, gene expression, SMAD factor partnership profiles, and transcription factor occupancy at target gene regulatory sequences.

## Materials and Methods

The list of all the reagents used during cell culture is given in **Table 1**, and the list of other materials is given in **Table 2**.

**Table 1.** List of reagents used in cell culture.

| Product | Provider | Reference | Stock Concentration (when relevant) | Solvent (when relevant) | Long-term Storage |
| --- | --- | --- | --- | --- | --- |
| ACTIVIN A (recombinant, mouse) | Cell Guidance System | GFM29 | 20 µg/ml | H2O + 0.1% BSA | -80°C |
| B27 | ThermoFisher | 17504-044 |  |  | -20°C |
| BMP-4 (recombinant, mouse) | R&D Systems | 5020-BP | 50 µg/ml | H2O + 0.1% BSA | -80°C |
| BSA 7.5 % | ThermoFisher | 15260-037 |  |  | 4°C |
| CHIR99021 | Cell Guidance System | SM13 | 3 mM | DMSO | -80°C |
| DMEM, high glucose, GlutaMAX™ Supplement, pyruvate | ThermoFisher | 31966021 |  |  | 4°C |
| DMEM/F-12, GlutaMAX™ Supplement | ThermoFisher | 31331028 |  |  | 4°C |
| DMSO | Euromedex | UD8050-B |  |  | 20°C |
| Fetal Bovine Serum, qualified | ThermoFisher | 10270106 |  |  | -20°C |
| FGF2 (recombinant, mouse) | Cell Guidance System | GFM12 | 10 µg/ml | H2O + 0.1% BSA | -80°C |
| Fibronectin from bovine plasma | Sigma-Aldrich | F1141 | 1mg/ml | PBS | 4°C |
| Gelatin from porcine skin | Sigma-Aldrich | G1890 | 0,15% | PBS | 4°C |
| H2O (UltraPure™ Dnase/Rnase-Free Distilled Water) | ThermoFisher | 10977035 |  |  | 4°C |
| KSR (KNOCKOUT™ Serum Replacement) | ThermoFisher | 10828010 |  |  | -20°C |
| L-Glutamine (200 mM) | ThermoFisher | 25030024 |  |  | -20°C |
| LDN193189 | Cell Guidance System | SM23 | 200 µM | H2O | -20°C |
| LIF (recombinant, mouse) | Cell Guidance System | GFM200 | 10 µg/ml (10<sup>6</sup> U/ml) | H2O + 0.1% BSA | -80°C |
| N2 | ThermoFisher | 17502-048 |  |  | -20°C |
| NeuroBasal | ThermoFisher | 21103-049 |  |  | 4°C |
| PBS (DPBS, no calcium, no magnesium) | ThermoFisher | 14190094 |  |  | 4°C |
| PD0325901 | Cell Guidance System | SM26 | 1 mM | DMSO | -80°C |
| Penicillin-Streptomycin (10,000 U/mL) | ThermoFisher | 15140122 |  |  | -20°C |
| Pluronic F127 | Sigma-Aldrich | P2443 | 1% | PBS | 4°C |
| Trypsin-EDTA (0.05%), phenol red | ThermoFisher | 25300054 |  |  | -20°C |
| WNT-3A (recombinant, mouse) | PeproTech | 315-20 | 100 µg/ml | H2O + 0.1% BSA | -80°C |

**Table 2.** List of reagents used before or after cell culture.

| Product | Provider | Reference |
| --- | --- | --- |
| Anti-Digoxigenin-AP Fab fragments | Sigma-Aldrich | 11093274910 |
| Blocking Reagent | Sigma-Aldrich | 46925300 |
| BSA (Bovine Serum Albumin) | Sigma-Aldrich | A3294 |
| Ethanol absolute | Sigma-Aldrich | 1,00983 |
| Fluoromount-G™ Mounting Medium | Thermofisher | 00-4958-02 |
| Formamide | Sigma-Aldrich | F9037 |
| Glutaraldehyde 25% | Sigma-Aldrich | G6257 |
| Héparine | Sigma-Aldrich | H3393 |
| Hoechst 33342 | Thermofisher | H3570 |
| Maleic acid | Sigma-Aldrich | M0375 |
| PFA (Paraformaldehyde) | Sigma-Aldrich | P6148 |
| Sylgard 184 Sicilone Elastomer Kit | DOW | 1673921 |
| Triton™ X-100 | Sigma-Aldrich | T8787 |
| Tween20 | Sigma-Aldrich | P7949 |
| Torula Yeast RNA | Sigma-Aldrich | 10109223001 |

### Basal medium

The basal medium used was N2B27, prepared as followed: mix 250ml DMEM/F12 with Glutamax, 250ml Neurobasal, 5ml B27 supplement, 2.5ml N2 supplement, 3.33ml 7.5% BSA solution, 5ml L-glutamine 200 mM and 5ml Penn/Strep 10 000 U/ml. The medium was then filtered using a 0.22 µm filtration unit, aliquoted in 50 ml conical tube and stored at -20C. Other constituents were added freshly the day of use.

### Routine mESC culture

ESCs were cultured on 0.15% gelatin coated tissue culture grade plates in 2i+LIF medium (N2B27 + 0.1 mM β-mercaptoethanol + 1000 U/ml LIF + 1µM PD0325901 + 3µM CHIR99021), in an incubator set at 37°C and 5% CO2. They were passaged every 2-3 days by dissociating them in 0.05% Trypsin for 5 min at 37°C. The trypsin was neutralized with DMEM + 10% FBS, the cells were resuspended in N2B27 and plated at around 1000 cells/cm².

The cell lines used are: HM1 WT (Selfridge et al., 1992) and E14Tg2a *Gsc*^GFP/+^ *Hex*^RedStar/+^(Villegas et al., 2013).

When the cell line is not mentioned, HM1 WT was used. When immunofluorescence of *Gsc*^GFP^ is shown, E14Tg2a *Gsc*^GFP/+^ *Hex*^RedStar/+^ was used.

### Production of micropatterned adhesive substrates

The micro-patterned substrates were produced using standard micro-contact printing approach. First, a mold with the desired pattern was produced by photo-lithography of SU8 resin on a silicon wafer. Uncured PDMS (Sylgard 184) was then poured on these molds to obtain a negative replica of the SU8 master, after curing at 65C for at least 1h. Cured PDMS was then peeled of the SU8 master and inked with a solution of fibronectin (50ug/ml in PBS, incubation 30 minutes at room temperature). The inked stamps were then washed twice with PBS and once with water and allowed to air dry for 10 minutes. During that time, PDMS-coated glass cover slips (obtained by spin coating PDMS on glass and curing) were activated in a UV-Ozone cleaner. The Fibronectin coated stamp and the activated coverslip were then put into contact, for a few seconds and separated. The glass cover slips were mounted in a magnetic chamlide. To prevent cell adhesion outside of the desired areas, stamped coverslips were then incubated in PBS + 1% Pluronic F127 for at least 30 minutes. After 3 washes with PBS, the micropatterned substrates were ready for cell seeding or could be stored at 4C for a future experiment.

### Differentiation of EpiLCs on micropatterns

ESCs cultures in 2i+LIF medium were harvested by trypsinization and seeded cells were first cultured for 24h on plates coated with fibronectin (15ug/ml for 30 min) in N2B27 supplemented with 0.1 mM β-mercaptoethanol and 1% KSR and optionally with 20ng/ml ACTIVIN and 12ng/ml FGF2. After 24h, cells were then dissociated using trypsin and seeded on micropatterned substrates at a density of 8000 cells per mm² , this high seeding density ensured to achieve a dense and uniform surface coverage. After 1h of incubation at 37°C, cells that did not attach were removed by gentle flushing and changing the culture medium and the remaining cells were allowed to form colonies for an additional 24h in 250µl of the same base medium. The medium was renewed every 24h during the course of the experiment. Colonies were stimulated by adding different combinations of factors to the medium. The stimulation was renewed after 24h, and colonies were cultured again in base medium.

### Immunostaining and microscopy

Cells were fixed in PBS + 4% paraformaldehyde at room temperature (RT) for 30 min, then washed three time with PBS for 5 min at RT, and blocked in a blocking solution (PBS + 0.1% Triton-X100 + 3% bovine serum albumin) for 1h at RT. They were then incubated overnight with the primary antibodies in the blocking solution (**Table 3**). The cells were then washed 3 times 5 min in PBS and once for 30 min with the blocking solution, incubated for 2 hours with a mix of secondary antibodies, wash three times in PBS (5 min + 15 min with 5 µM Hoechst + 5 min) and mounted in Fluoromount-G. Fluorescent images of the colonies were acquired with an Olympus IX81 inverted microscope equipped with a Yokogawa CSU-X1 spinning disk head. Radial profiles and co-expression analyses of fluorescence were extracted from maximum intensity projection using a custom python script.

**Table 3.**
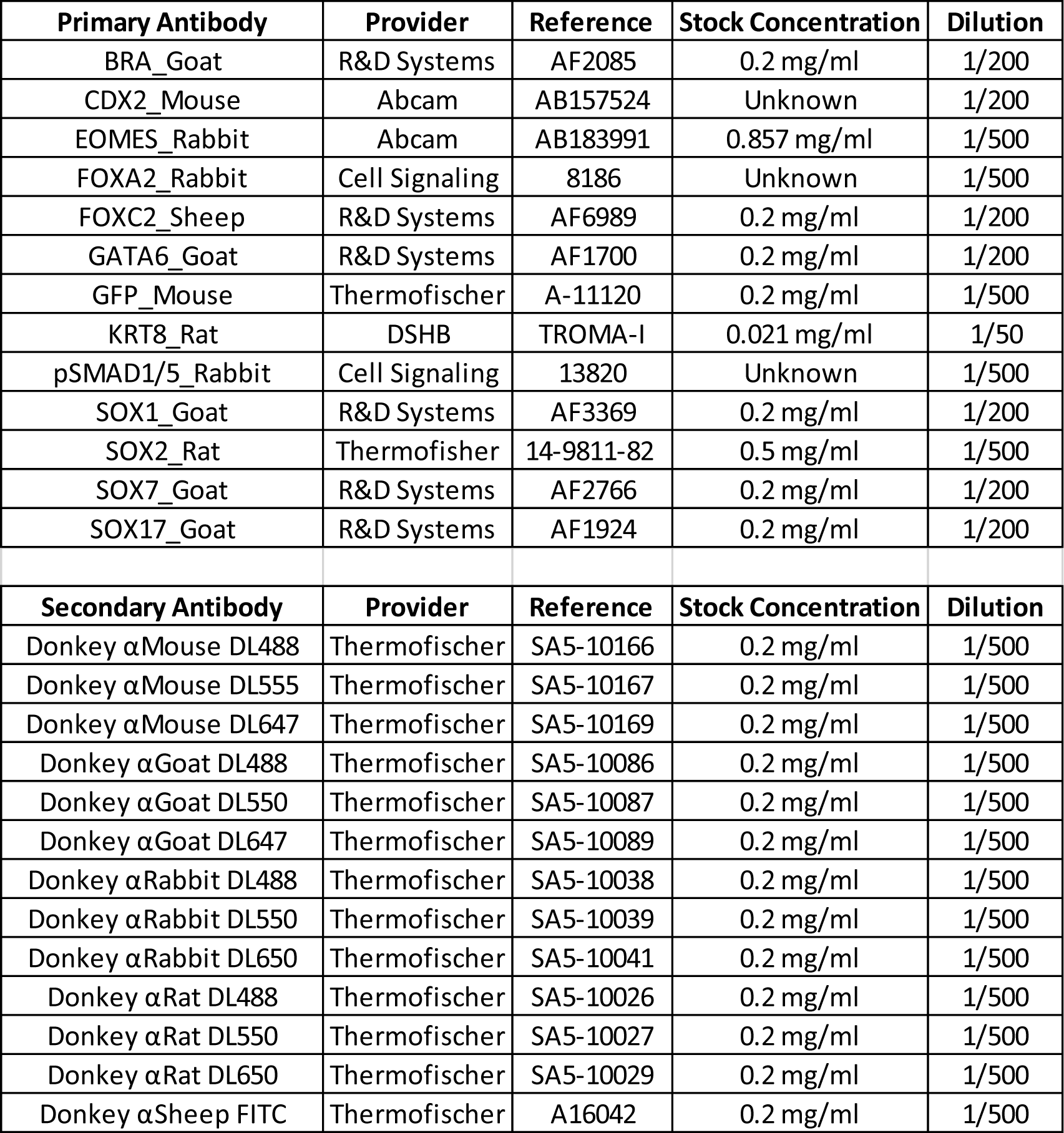
List of antibodies used for IF.

### Quantitative analysis of gene expression

Cell total RNA was extracted using the RNANucleoSpin RNA extraction kit (Macherey-Nagel). After the cell lysis in the RA1 solution + 1% β-mercaptoethanol, the samples were kept at - 80°C until the further steps. RNA quantification was performed using a Nanodrop. Between 25 and 200ng of total RNA were then reverse transcribed (RT) using the Superscript IV VILO reverse-transcriptase (Thermo Fisher Scientific) and random *Hex*amers according to manufacturer’s instructions. Gene expression was quantified by quantitative PCR on a LightCycler 480 Instrument II (Roche) using 1:50 of the reverse-transcribed RNA sample mixed with a specific primer pair (1uM final each, sequence in **Table 4**) and 2X KAPA SYBR FAST qPCR Master Mix (Roche) according to manufacturer’s conditions. For each sample, technical duplicates samples were analyzed. For each gene, the quality of the amplification was tested by quantifying its expression in a serially-diluted pool of all RT (1:10 to 1:320) to quantify amplification efficiency and target mRNA concentration relative to the RT pool in each sample. A custom R script was used to compute the following steps. We computed for each sample and gene a relative gene expression value with respect to the mean expression of the gene in the experiment (obtained by pooling all samples belonging to the same experiment) and normalized by the expression of the GAPDH gene in the sample. This value was log2- transformed, then centered and reduced with respect to the expression value of the gene in all samples considered in order to compute and display the expression matrix and the principal component analysis of this matrix. When biological replicates were performed, the 95% confidence ellipse was computed using the *plot.PCA* function from the *FactoMineR* package.

**Table 4.**
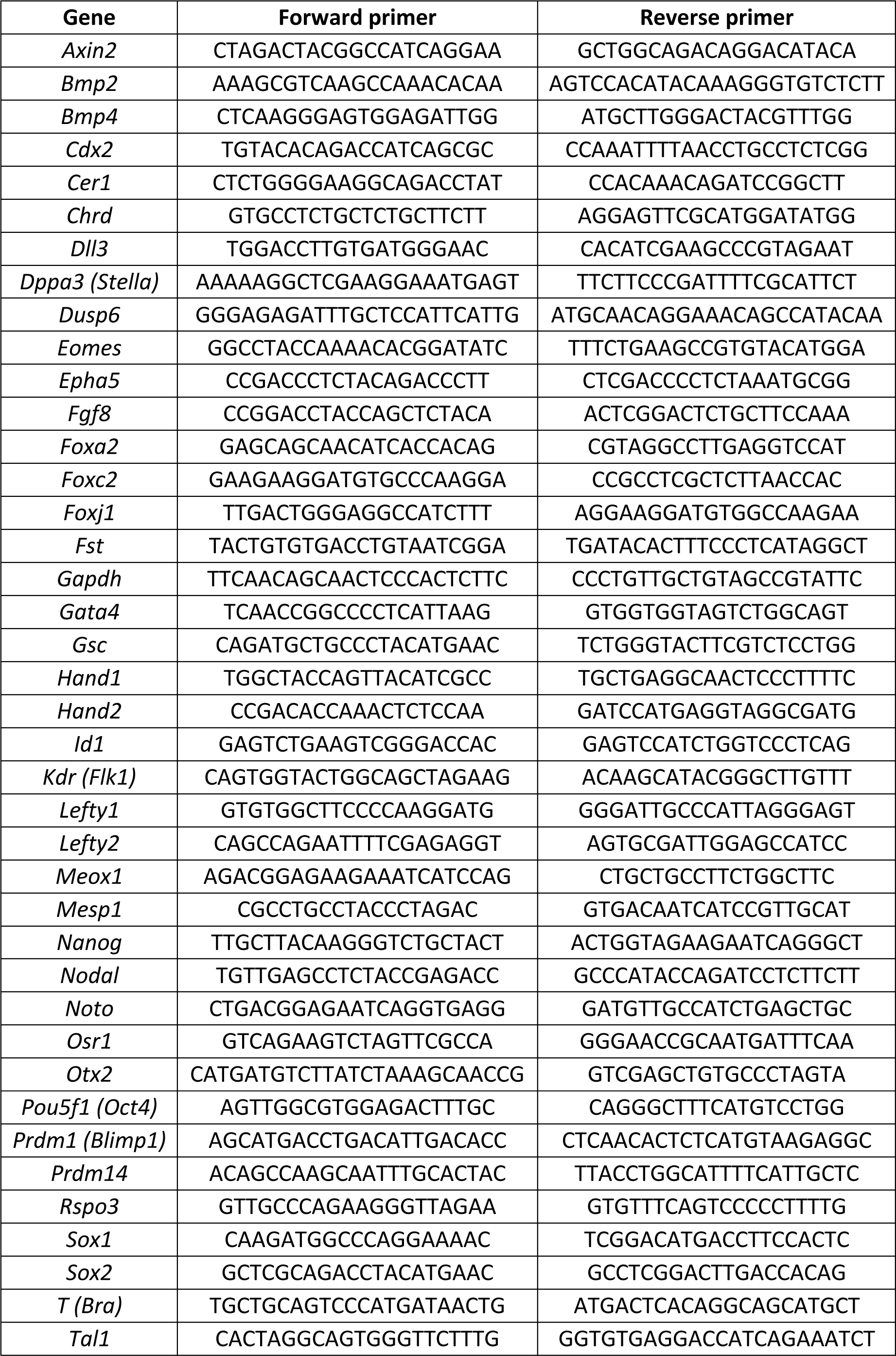
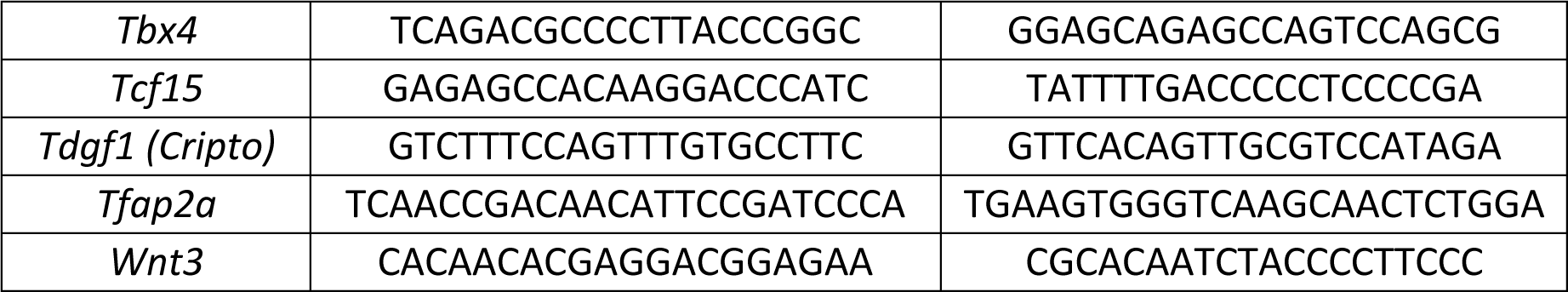
List of primers used for RT-qPCR.

### In-situ Hybridization (ISH)

List of the solution used during ISH:

PBT (Store at RT): PBS + Tween20 0.1% Hybridization solution (Store at -20°C):

- 25 ml of Formamide

- 12,5 ml of SSC 20X

- 5 ml of Dextran Sulfate 50%

- 250 µl of Yeast RNA 10 mg/ml

- 250 µl of Heparin 10 mg/ml

- 500 µl of Tween20 10%

- 6,5 ml of H20

Wash1 (Store at -20°C): 50% Formamide, 2X SSC, 0,1% Tween20, in H20 Wash2 (Store at -20°C): 2X SSC, 0,1% Tween20, in H20 Wash3 (Store at -20°C): 1X SSC, 0,1% Tween20, in H20 BBR (Store at -20°C):

- 20 ml of Maleate solution 250mM pH7.2

- 1,5 ml of NaCl 5M

- 500 µL of Tween20 10%

- 28ml of H20

- 2,5g of Blocking Reagent (to be added last, slowly at 60°C with agitation) NTMT (prepare freshly the day of use):

- 1 ml of Tris 1M ph 9,5

- 200 µl of NaCL 5M

- 500 µl of MgCl2 1M

- 100 µL of Tween20 10%

- 8.2ml of H20

(Day0) Cells were fixed in PBS + 4% paraformaldehyde at RT for 4 hours, then washed three time with PBT for 5 min, and dehydrated in Ethanol (25 -> 50% -> 75% -> 100%, 5 min each) and stored at -20°C until use.

(Day1) Cells were rehydrated from ethanol to PBT (75 -> 50% -> 25% -> PBT, 5 min each), and washed three more times with PBT for 5 min. Cell were post-fixed for 20 min in PBS + PFA 4% + 0,2% Glutaraldehyde and washed three times with PBT for 5 min at RT. Cells were put in Hybridization buffer for 1h at 67° and then overnight at 67°C in Hybridization buffer + 1 µg of DIG-labelled RNA probe (Day2) The cells were washed 5 min at 67°C in Wash1, 2x15 min at 67°C in Wash1, 15 min in Wash 2 (RT from now), 2x15 min in Wash3, 3x5min in PBT, 5 min in BBR. Cells were blocked in BBR + 10% FBS for 1h and incubated in BBR + 10% FBS + 1/2000 AP-anti-DIG antibody for 4h. The cells were washed three times with PBT for 5 min and stored for the night in PBT at 4°C (Day3) Fresh NTMT was prepared. The cells were washed two times with PBT for 5 min, in NTMT for 5 min and in NTMT for 30 min. During this time, a revelation solution of 3 ml of NTMT + 13,5 µl of NBT + 10,5 µl of BCIP was prepared and kept away from light. The cells were then put in the revelation solution and frequently monitored under a loop. Once the staining was sufficient, the cells were washed three times with PBT for 5 min. Once all the cells were wash, the coverslip was mounted in 80% Glycerol and stored at 4°C until the imaging on a Leica microscope DM5000B.

### 10X Single cell RNA sequencing

The colonies were dissociated in 0.05% Trypsin for 5 min at 37°C. The trypsin was neutralized with DMEM + 10% FBS, the cells were resuspended twice in PBS + 0.04% BSA. scRNA-seq libraries were subsequently generated using the 10x Genomics Chromium system and samples were sequenced according to the manufacturer’s instructions on an Illumina NextSeq 500. The demultiplexing and the count matrices for each time point were obtained using Cellranger

#### 2.1.1. The following analyses were performed using the *Seurat* package in R

#### Quality control and normalization

Cells with less than 10 000 UMI counts, with more than 40 000 UMIs or with more than 5% mitochondrial genes were excluded. Cell counts were normalized using the function ‘NormalizeData’ to have a count per cell of 20 000, before performing a log-transformation. Variables features were selected using the function ‘FindVariableFeatures’. 3000 were kept prior to the integration of all the time points while 2000 were kept for the individual analyses of time points.

#### Principal Component Analysis (PCA)

Prior to PCA, the log-transformed data where centered and reduced using the function ‘*ScaleData*’. The PCA was then computed with the scale data of the variable features.

#### Integration using Mutual Nearest Neighbors (MNN)

The function ‘*RunFastMNN*’ was used to integrate and apply batch correction when different datasets were analysed together. For the integration of the 5 m2Dgas time points, 40 principal components (PC) and 5 MNN were used as parameters, while for the comparison of one m2Dgas time point to one embryonic time point, 50 PCs, 10 MNNs and the embryonic dataset as reference were used.

#### Clustering and UMAP

The clustering was performed on the individual time points. For t00 and t06, no clustering was performed and they were both annotated as Epi. The clustering was performed using the functions ‘*FindNeighbors*’ and ‘*FindClusters*’. For the individual time points, the number of neighors (k.param) was set at 10, and the 10, 30 and 30 PC were taken for respectively t24, t48 and t72. The same parameters were used to compute the UMAP with the function ‘RunUMAP’. To generate clustering trees (Fig. S3A and S4A), the clustering was performed using different resolutions and the package ‘clustree’ was used to generate the plots.

For the comparison between m2Dgas and embryo, the batch-corrected PCA space from the ‘*RunFastMNN*’ results were used with the default parameters of the UMAP function, except for the number of PCs. The first 30 PCs were used for t00vsE5.5 and t24vsE6.5, while the first 50 PCs were used for t48vsE7.5 and t72vsE8.25.

#### Differentially expressed genes (DEG) analysis

The function ‘*FindAllMarkers*’ was used, with default parameters except for the parameter ‘*logfc.threshold*’ (0.2 for the figure S1D and S2B-C, 0.1 for the figures S4B and S4F). For the gene ontology analysis relative to the figure S2C, the libraries ‘mgsa’ and ‘topGO’ were used. The enrichment test was performed using the function ‘runTest’ with the ontology ‘Biological Process’ on the genes that had p_val_adj<0.01.

## Data availability

The single cell RNA sequencing data can be found on Gene Expression Omnibus under the accession number GSE276079.

## Authors contributions

BS and JC conceived the research, obtained funding and oversaw the experiments and data analysis. GS and JLP carried out the experiments and analyzed the data. GS, BS and JC wrote the manuscript.

## Competing interests

The authors declare no competing interests.

## Acknowledgements

We thank Fanny Coulpier from the genomic facility at ENS, for her help in the production and analysis of the scRNAseq data, and Hervé Isambert for discussions.

We also acknowledge the ImagoSeine core facility of Institut Jacques Monod, member of France-Bioimaging (ANR-10-INBS-04) and IBISA, with the support of Labex “Who Am I”, INSERM Plan Cancer, Région Ile-de-France and Fondation Bettencourt Schueller.

This work was supported by CNRS and by grants from INCa (2014-1-PLBIO-01), HFSP (CDA00063/2015-C), Ligue Contre le Cancer (RS23/75-86) and ANR (ANR-15-CE13-0007-01 and ANR-23-CE13-0018-03). GS was supported by Université Paris-Cité (ED562).

## Notes

### Competing Interest Statement

The authors have declared no competing interest.

