## Supplementary Figures for "BMP signaling modulations control primitive streak patterning"

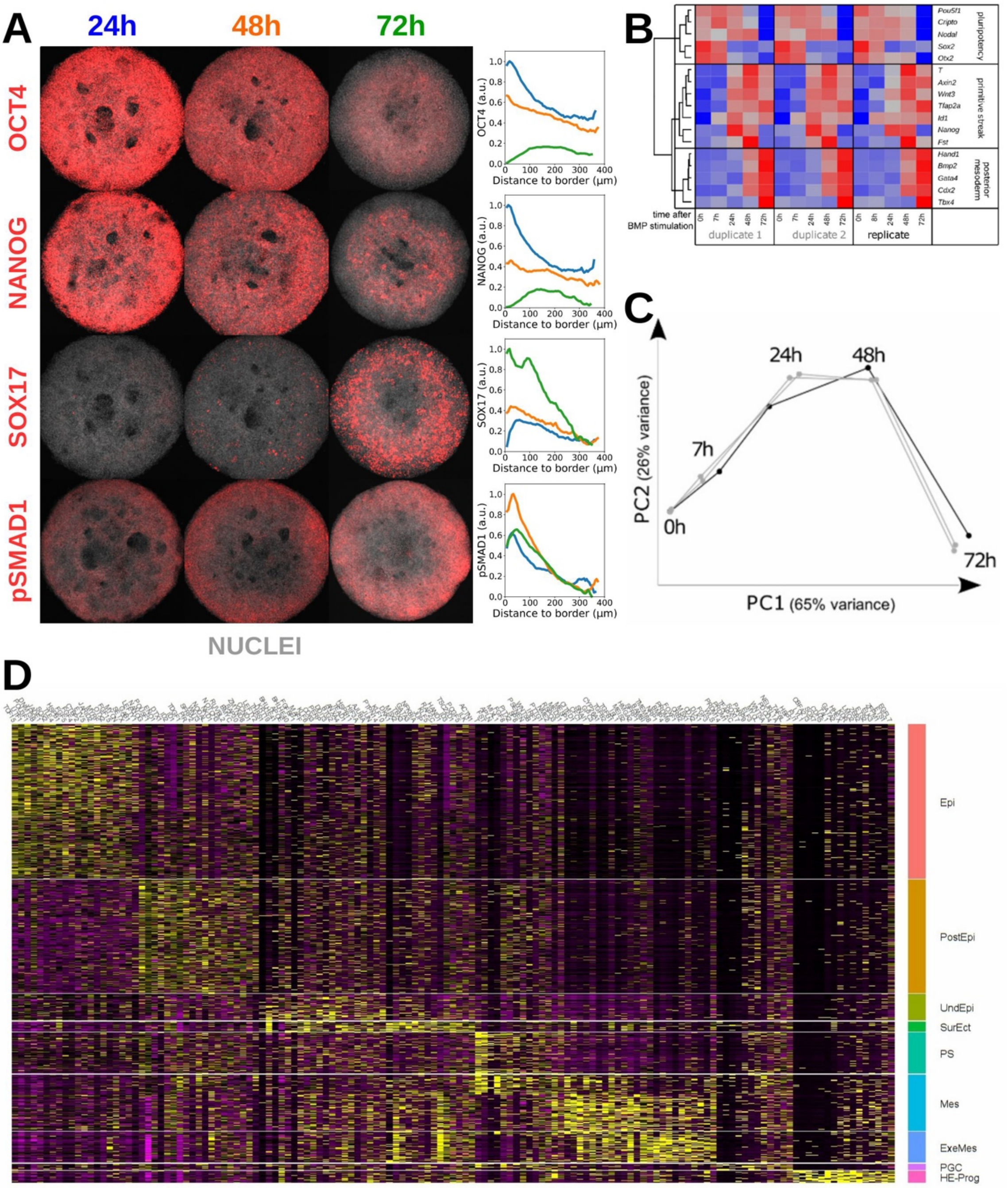


***Figure S1 (Supplementary Data for Figure 1)***

***(A)*** *Maximum intensity projections of immunostained m2Dgas (700 µm) 24, 48 or 72 hours after the start of BMP4 stimulation.* *Fluorescence intensity levels normalized by Hoechst and averaged along the colony radii (n=3 for each curve)*

***(B)*** *Gene expression matrix of multiple replicates obtained by qPCR 0/7/24/48/72h after BMP stimulation. Duplicates 1&2 belong to the same experiment as the scRNAseq. The other replicate belongs to an independent experiment. Genes were clustered according to the similarity of their expression. This panel was already published in (Plouhinec et al., 2022)*

***(C)*** *Temporal trajectory of the replicates along the first 2 principal components from the data of (C)*

***(D)*** *Heatmap of markers (transcription factors and known targets of signaling pathways) used to annotate the cell identities present at the different time points.*

*Epi: Epiblast, PostEpi: Posterior Epiblast, UndEpi: Undifferentiated Epiblast, SurEct: Surface Ectoderm, Mes: Mesoderm, ExeMes: Extra-embryonic mesoderm, PGC: Primordial Germ Cells, HE-Prog: Haematho-Endothelial Progenitor.*

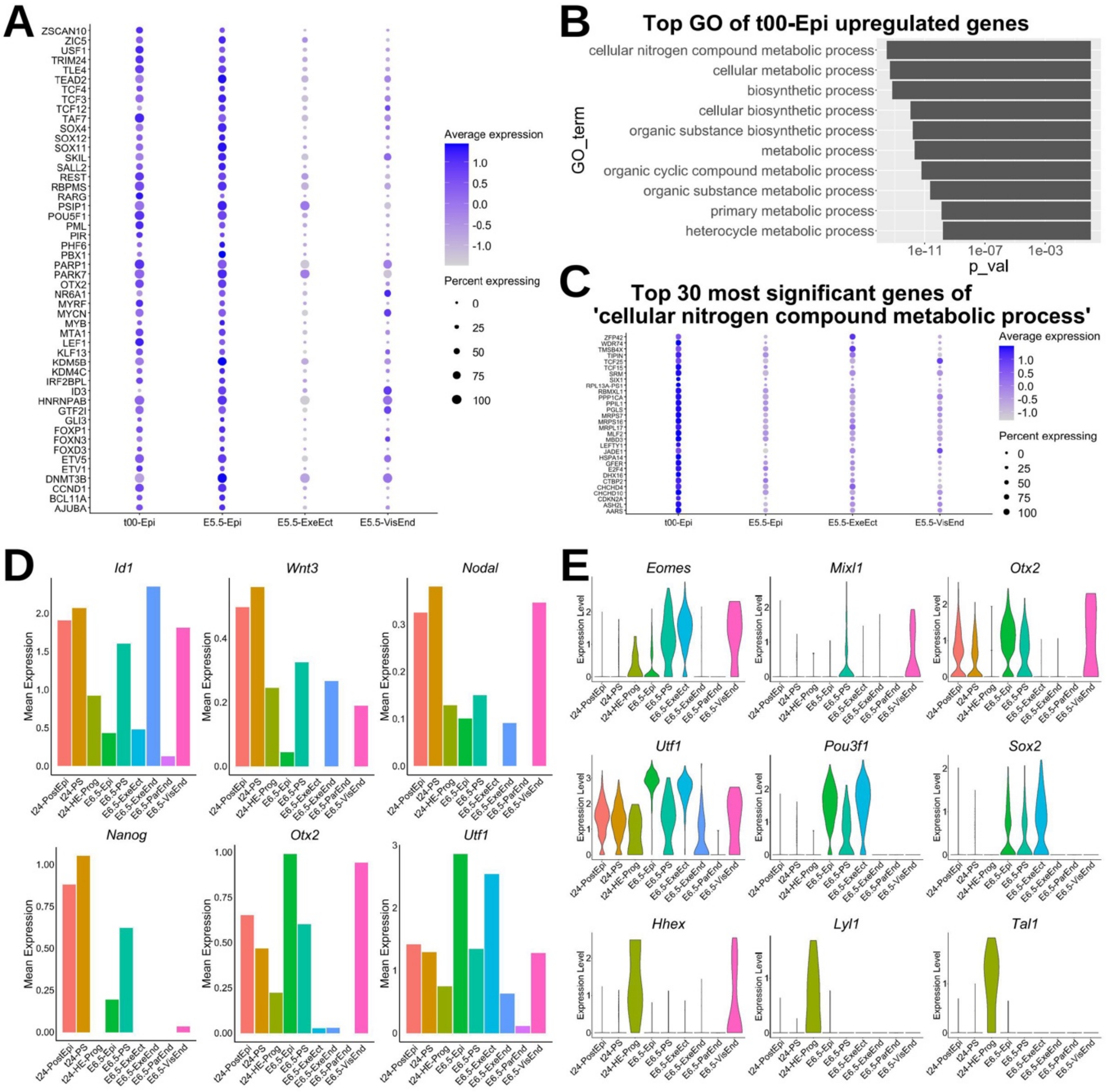


***Figure S2 (Supplementary Data for Figure 2)***

***(A)*** *Dot plot of the expression of the 50 most upregulated genes of E5.5-Epi versus E5.5-ExeEct and E5.5-VisEnd.*

***(B)*** *Gene ontology analysis of the upregulated genes of t00-Epi versus E5.5-Epi identifies differences in metabolic processes.*

***(C)*** *Dot plot of the expression of the 30 most significant genes from the gene ontology term "cellular nitrogen compound metabolic process", relative to* ***(B).***

***(D)*** *Bar plots of the mean expression of genes linked to posterior epiblast formation.*

***(E)*** *Violin plots of key genes involved in primitive streak formation, pluripotency and HE-prog formation.*

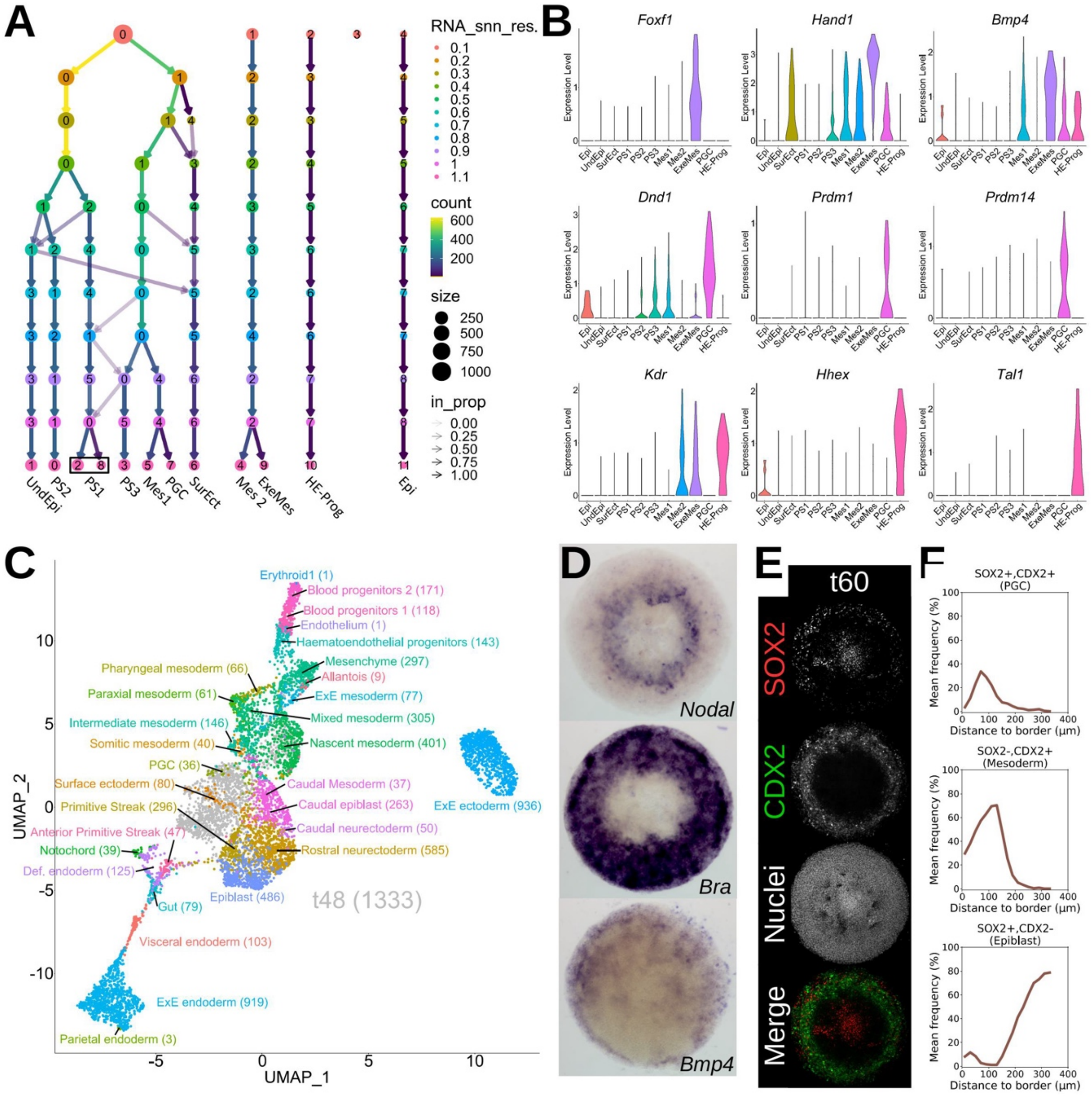


***Figure S3 (Supplementary Data for Figure 3)***
***(A)*** *Clustering tree relative to Fig. 3A. As the resolution increases, the number of clusters also increases. We have used a resolution of 1.1 as this allows the ExeMes and PGC clusters to appear.*
***(B)*** *Violin plots of key genes involved in ExeMes, PGCs and HE-Prog formation.*
***(C)*** *UMAP representation of the integrated dataset for t=48h m2Dgas cells and E7.5 mouse embryo cells (Pijuan-Sala et al., 2019). Only the E7.5 clusters are colored and labelled.*
***(D)*** *In situ hybridization of 700 µm m2Dgas at t=48h for Nodal, Bra (T) and Bmp4.*
***(E)*** *Maximum intensity projections of 700 µm m2Dgas immunostained for SOX2 and CDX2 at t=60h.*
***(F)*** *Single cell co-expression analysis relative to* ***(E)****, and frequencies averaged along the colony radii (n=6).*

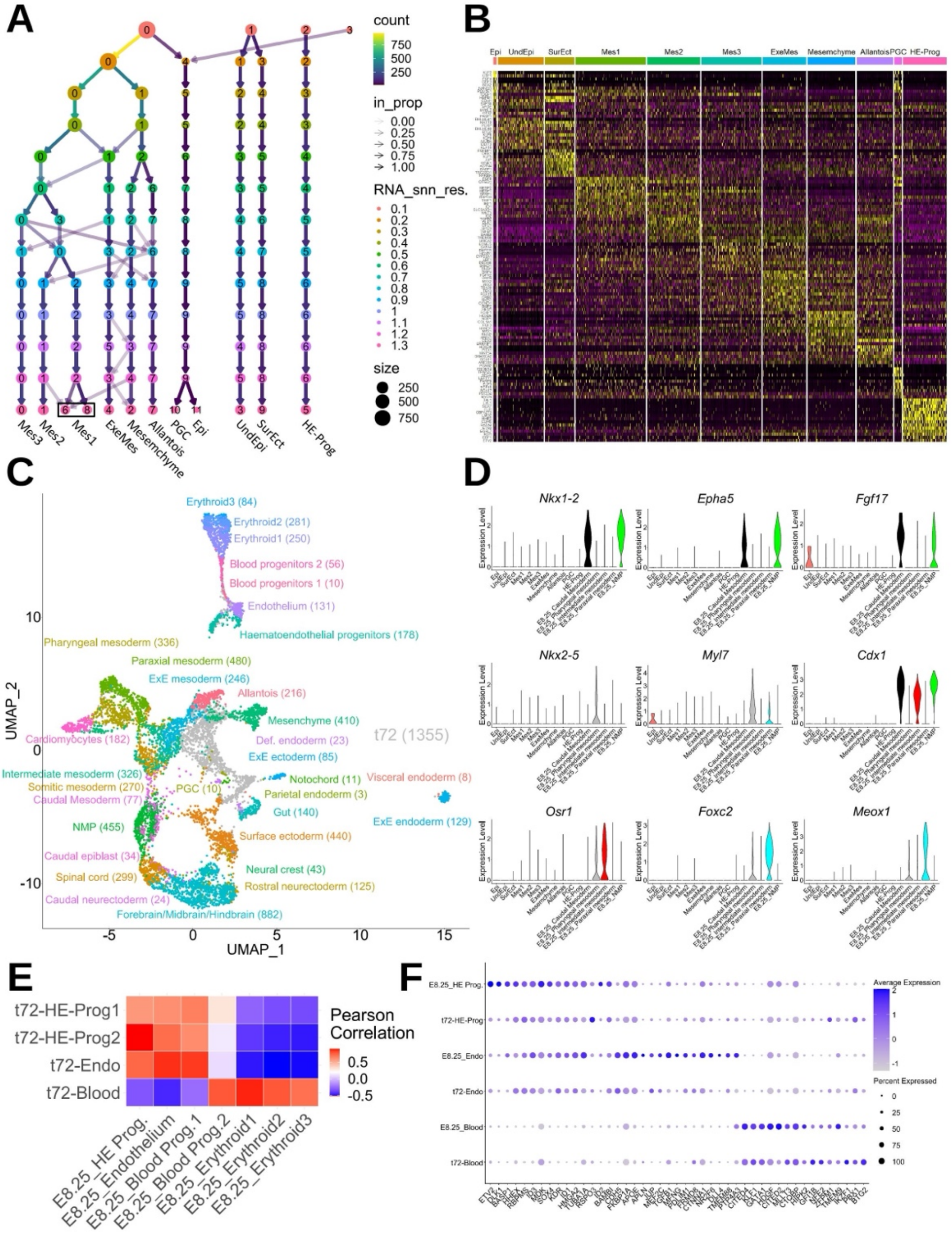


***Figure S4 (Supplementary Data for Figure 4)***

***(A)*** *Clustering tree relative to Fig. 4A. As the resolution increases, the number of clusters also increases. We used a resolution of 1.3 as it allows the identification of a PGC cluster.*

***(B)*** *Heatmap of markers (transcription factors and known targets of signaling pathways) used to annotate the cell identities present at t=72h.*

***(C)*** *UMAP representation of the integrated dataset for t72-m2Dgas cells and E8.25 mouse embryo cells (Pijuan-Sala et al., 2019). Only the E8.25 clusters are colored and labelled.*

***(D)*** *Violin plots of key genes involved in embryonic mesoderm specification.*

***(E)*** *Mean Pearson correlation in the batch-corrected PCA space between the subclusters of t72-HE-Prog and their E8.25 embryonic counterparts.*

***(F)*** *Dot plot showing the expression of the top 15 markers (transcription factors only) for the cell types shown in (E). The cell types Blood Prog.1, Blood Prog.2, Erythroid1, Erythroid2, Erythroid3 were grouped together and called Blood.*

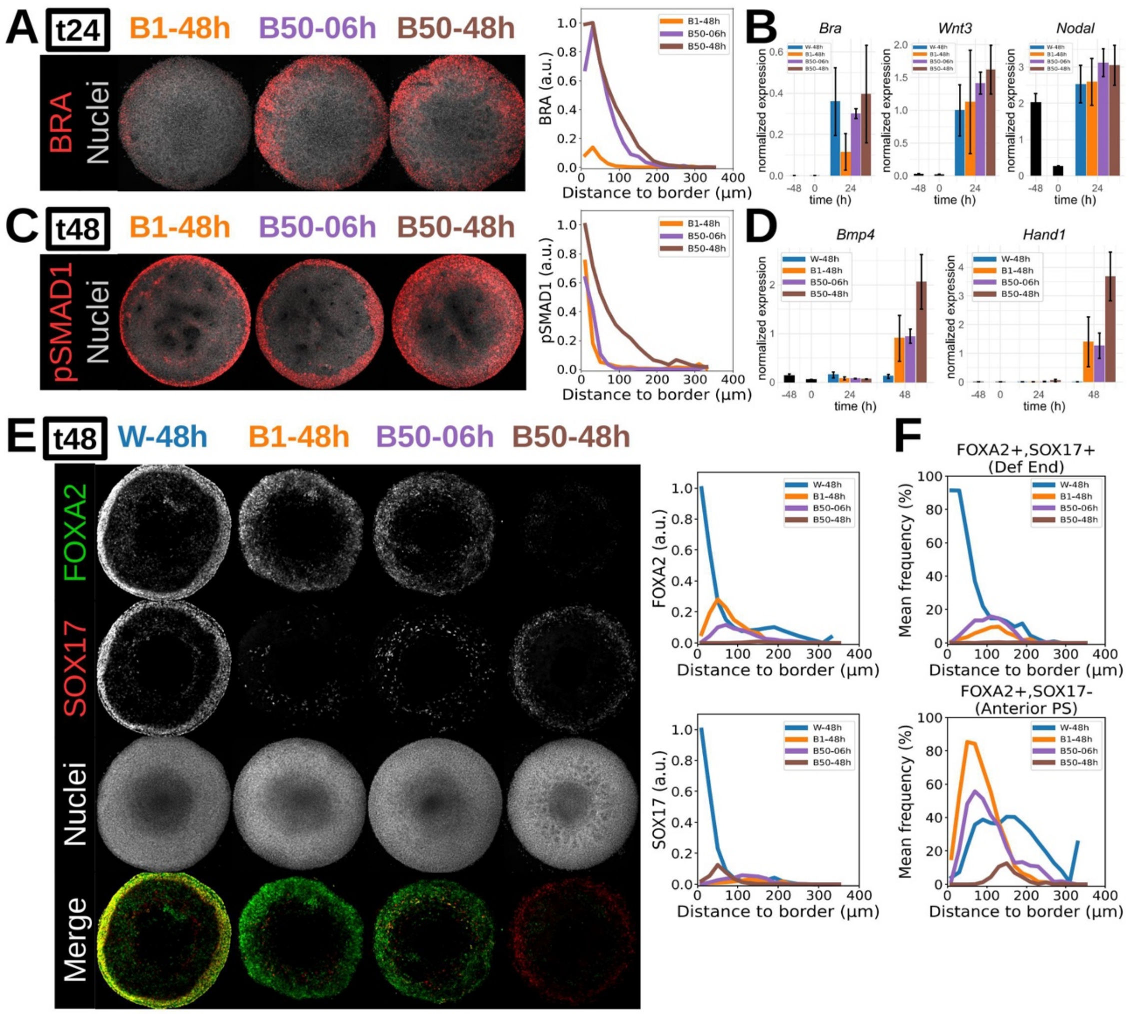


***Figure S5 (Supplementary Data for Figure 5)***

***(A)*** *Maximum intensity projections of immunostained m2Dgas (700 mm) for BRA after 24h of differentiation, following the different protocols. Fluorescence intensity levels normalized by Hoechst and averaged along the colony radii (n=3 for each curve).*

***(B)*** *Gene expression measurements obtained by RT-qPCR of whole colonies (mean ± SEM, n=3).*

***(C)*** *Maximum intensity projections of immunostained m2Dgas (700 mm) for pSMAD1 after 24h of differentiation, following the different protocols. Fluorescence intensity levels normalized by Hoechst and averaged along the colony radii (n=4 for each curve).*

***(D)*** *Gene expression obtained via RT-qPCR of whole colonies (mean ± SEM, n=3).*

***(E)*** *Maximum intensity projections of immunostained m2Dgas (700 mm) for FOXA2 and SOX17 after 48h of differentiation, following the different protocols. Fluorescence intensity levels normalized by Hoechst and averaged along the colony radii (n=4 for each curve).*

***(F)*** *Single cell co-expression analysis relative to (E), and frequencies averaged along the colony radii (n=4).*

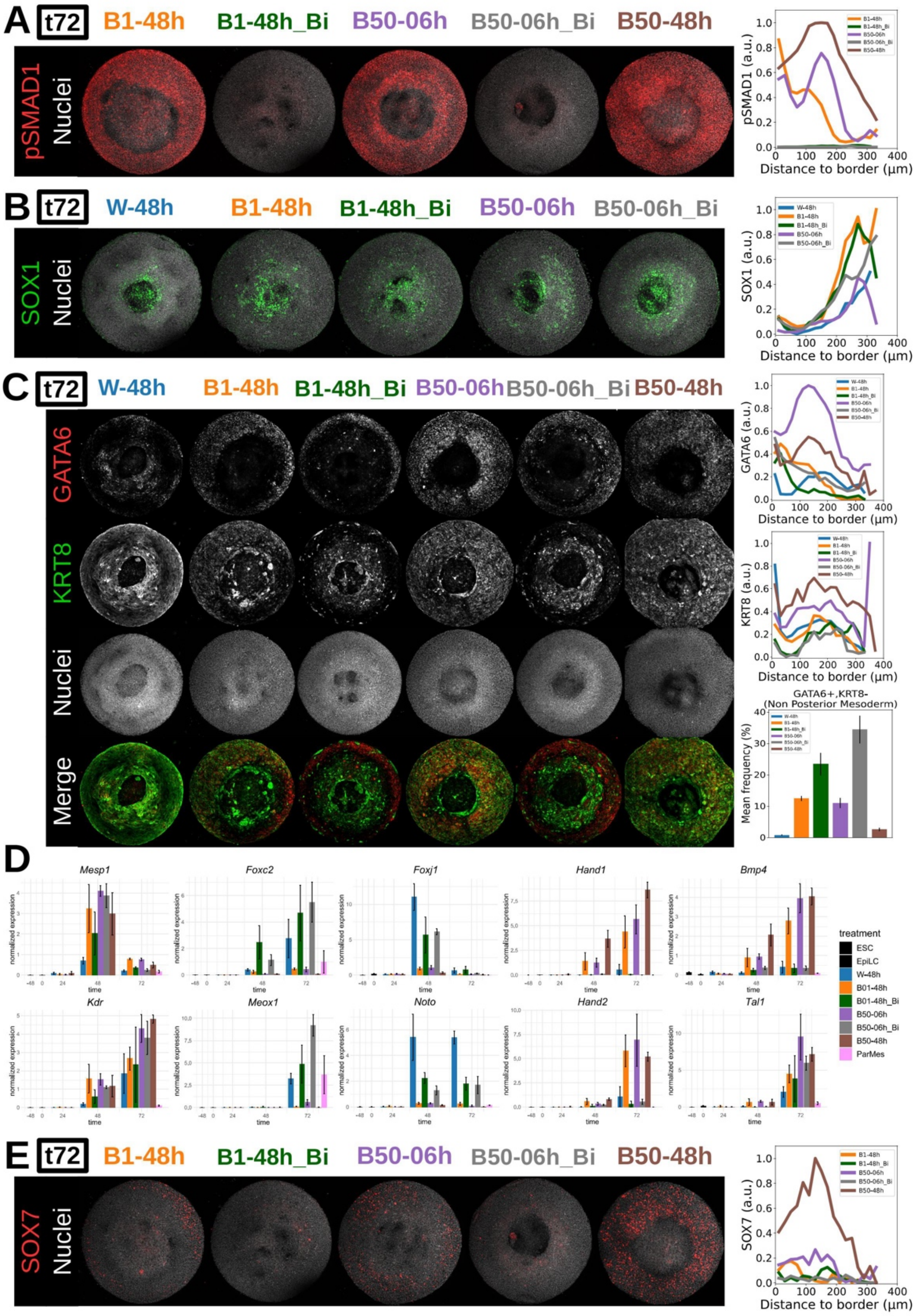


***Figure S6 (Supplementary Data for Figure 6)***

***(A, B, D, E)*** *Maximum intensity projections of immunostained m2Dgas (700 mm) after 72h of differentiation in the different culture conditions. Fluorescence intensity levels were normalized by Hoechst and averaged along the colony radii (n=4 for each curve).*

***(A)*** *BMP signaling inhibition results in the loss of pSMAD1 staining.*

***(B)*** *SOX1 expression (neural ectoderm) remains restricted to the center of the colonies after BMP signaling inhibition.*

***(C)*** *Non-posterior mesoderm was defined as GATA6+KRT8-, and BMP signaling inhibition leads to an increase in the frequency of GATA6+KRT8- cells in the colonies.*

***(D)*** *Gene expression obtained by RT-qPCR of whole colonies (mean ± SEM, n=3).*

***(E)*** *Inhibition of BMP signaling results in loss of SOX7 expression, an HE progenitor marker, in the outer ring of the colonies.*
